# A stage-resolved map of dynamic septin interactions required for infection by the rice blast fungus

**DOI:** 10.64898/2026.04.02.716073

**Authors:** Iris Eisermann, Neha Sahu, Marisela Garduño Rosales, Paul Derbyshire, Frank L.H. Menke, Weibin Ma, Nicholas J. Talbot

**Affiliations:** The Sainsbury Laboratory, University of East Anglia, Norwich Research Park, Norwich NR4 7UH, UK

**Keywords:** septins, interactome, cytoskeletal organisation, cell polarity, membrane remodelling, protein-protein interactions, appressorium development, fungal pathogenesis, *Magnaporthe oryzae*

## Abstract

Septin GTPases are essential cytoskeletal regulators that organize membranes and scaffold protein complexes to control cytokinesis, polarity, and morphogenesis. How septins execute these functions remains poorly understood, and comprehensive, stage-resolved interaction maps are lacking. Here, we define a quantitative, time-resolved septin interactome in the rice blast fungus *Magnaporthe oryzae* using immunoprecipitation coupled to mass spectrometry. We map more than 350 interactors of septins Sep3, Sep4, Sep5 and Sep6, revealing a dynamic network required for appressorium-mediated plant infection. Beyond canonical roles in cytoskeletal organisation and polarity, septins associate with proteins linked to membrane remodelling, metabolism, and virulence, deployed during host invasion. Integration with ultra-high-throughput yeast two-hybrid analysis defines a high-confidence septin interactome and identifies previously uncharacterised factors, including Msi1, a BAR domain protein required for invasive growth. Together, these findings establish septins as dynamic organisers of infection-related processes and provide a framework for understanding how cytoskeletal scaffolds coordinate fungal pathogenesis.

## Introduction

Septins are a conserved family of GTP-binding cytoskeletal proteins that assemble into hetero-oligomeric complexes and higher-order structures at membranes, where they function as scaffolds and diffusion barriers.^1,2^ Septin polymers interface with actin and microtubules and are essential for cytokinesis, cell polarity establishment and membrane remodelling across eukaryotes.^1,3,4^ Despite these fundamental roles, how septins execute their diverse cellular functions remains poorly understood, particularly at the level of their dynamic protein interaction networks.

Septins are widely conserved among opisthokonts and many other eukaryotes but notably absent from land plants.^1^ At the cell cortex, septins assemble into higher-order structures, including bars, rods, rings, gauzes, and collar-like arrays that shape membrane organisation, promote substrate interactions, and enable symmetry-breaking during development and host-pathogen interactions.^5,6^ In humans septin dysfunction is linked to cancer, neurodevelopmental and neurodegenerative disorders, thrombocytopenia and immune defects, and also contribute to host defence by restricting invading pathogens.^1,3,7^ Much of our mechanistic understanding of septin assembly derives from budding yeast, where septins form ordered cortical structures, including rings and hourglass assemblies at the mother-bud neck, that act as paradigmatic organisers of polarity and cytokinesis.^8,9^ These studies have established key principles of septin assembly but provide limited insight into how septins are dynamically integrated with broader cellular processes.

A central unresolved question is how septins are connected to organelles, metabolic pathways, and signalling networks that coordinate morphogenesis, stress responses, and infection-related development. To date, most studies have focused on static localisation patterns or a limited number of interactors, leaving the composition, temporal remodelling, and functional organisation of septin-associated protein networks largely unexplored.^3,9–11^ As a result, the extent to which septins act as systems-level organisers to integrate cytoskeletal architecture with cellular physiology remains unclear across eukaryotes.

Quantitative interactome profiling approaches such as co-immunoprecipitation coupled to mass spectrometry (Co-IP-MS), provide a powerful framework to address this gap by capturing assembly and remodelling of protein complexes across defined developmental states.^12,13^ When combined with parallelised experimental designs and rigorous statistical analysis, these approaches enable temporal changes in protein interactions to be resolved at systems scale.^14,15^ Complementary high-throughput yeast two hybrid (Y2H) screening provides an orthogonal view of direct binary interactions, allowing direct binding partners to be distinguished from co-complex associations and redefining network architecture at single-protein resolution.^16–20^ Integrating these datasets with network-based analyses enables reconstruction of dynamic interaction modules and offers a general framework to dissect how septin scaffolds are functionally wired within the cell.^21,22^

Septins are critical for virulence in both human and plant pathogenic fungi, where their perturbation leads to severe defects in infection-related development.^23–25^ In the rice blast fungus *Magnaporthe oryzae*, for example, which is a major threat to global food security, a heteromeric septin ring composed of Sep3, Sep4, Sep5 and Sep6 assembles at the appressorium pore.^24^ Septins scaffold a toroidal F-actin network, reinforce cortical rigidity, and enable repolarisation required for host invasion.^26,27^ Loss of any core septin disrupts appressorium function and severely reduces virulence. Despite these insights, the composition and temporal organisation of septin-associated protein networks during infection are poorly defined beyond a small number of characterised components.^24,27,28^

In this study, we performed time-resolved co-immunoprecipitation-mass spectrometry on four *M. oryzae* septins (Sep3, Sep4, Sep5, Sep6), during vegetative growth and appressorium development. We identified 1179 putative septin interactors, including a high-confidence interactome of 349 proteins, revealing extensive temporal remodelling of septin-associated networks during infection-related development. Complementary high-throughput yeast two-hybrid screening provided orthogonal support for these interactions defining a subset of *Mangnaporthe* septin interactor (Msi) proteins selected for functional analysis. Among these we identify Msi1 as a septin-associated BAR domain protein required for invasive growth and virulence. Together, our findings reveal how dynamic septin assemblies organise cellular architecture and infection programmes in a pathogenic fungus.

## RESULTS

### Time-resolved septin interactome analysis reveals distinct septin-associated networks during infection cell development by *Magnaporthe oryzae*

The four septins Sep3, Sep4, Sep5, and Sep6 assemble into hetero-oligomeric structures throughout appressorium development, as visualised by super-resolution microscopy (Figure 1). At early time points (0-2 h post-inoculation) septins displayed distinct localisation patterns: Sep3 formed bar- and tubule-like structures and puncta; Sep4 and Sep5 localised to septa, cytoplasmic filaments, and regions of positive membrane curvature at the plasma membrane; while Sep6 appeared as puncta and short filaments within the conidium and sub-apical to the Spitzenkörper at germ-tube tips. From 4 h onward, septins progressively co-assemble into hetero-oligomeric structures, initially forming a disc at the appressorium base (4-8 h) that matures into a ∼5 µm basal ring by 16-24 h (Figure 1A). Structured illumination microscopy of dual-labelled Sep4-GFP/Sep5-TagRFP strains revealed co-assembly of Sep4 and Sep5 into an interwoven, mesh-like network at the appressorium base (Figure 1B). This organisation is consistent with previous reports^24,29^ and supports a role for septins as a dynamic scaffold within infection structures.

**Figure 1.**
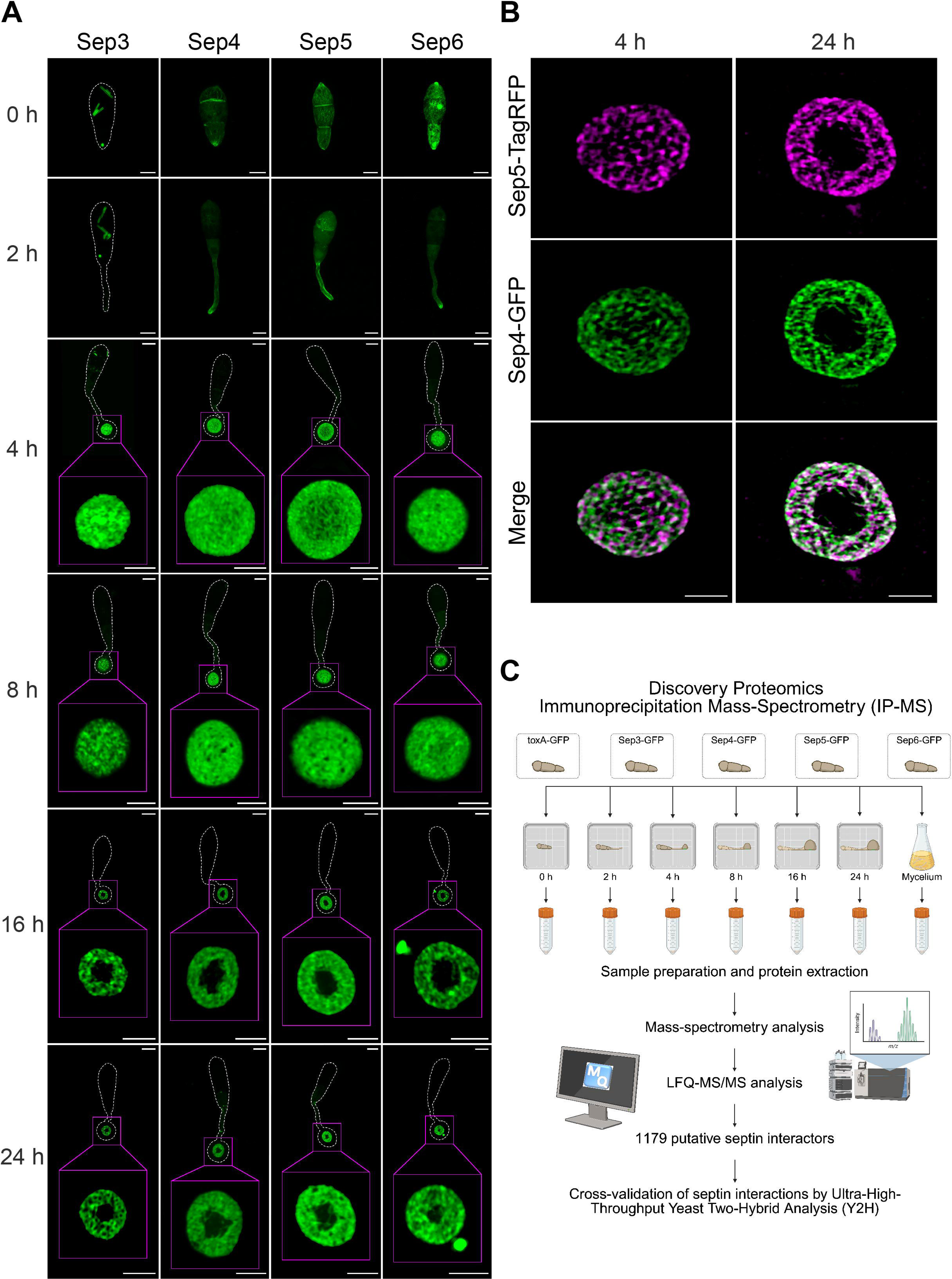
Septin localisation and strategy to define temporal dynamics of septin-dependent organisation of appressorium development in *M. oryzae*. (A) Confocal laser scanning microscopy combined with Airyscan super-resolution imaging reveals the dynamic localisation of the four core septins (Sep3-GFP, Sep4-GFP, Sep5-GFP and Sep6-GFP) at key stages of appressorium development (0, 2, 4, 8, 16, and 24 h post inoculation). Scale bars: 5 μm (whole-structural views) and 2 μm (magnified insets; outlined in magenta). (B) Structured Illumination Microscopy (SIM) provides super-resolution visualisation of septin hetero-polymers, captured as a disc-like structure at 4 h and a mature ring at 24 h at the base of the appressorium, using dual-labelled Sep4-GFP/Sep5-TagRFP strain. Scale bar: 2 μm. (C) Schematic overview of discovery proteomics workflow used to identify septin interactors during appressorium development and vegetative growth (mycelium). Comparative immunoprecipitation mass spectrometry (IP-MS) was performed on strains expressing GFP-tagged Sep3, Sep4, Sep5, and Sep6, or a toxA-GFP control across developmental stages (0, 2, 4, 8, 16, 24 h post inoculation; 48 h for mycelium). Analysis of three biological replicates by MaxQuant identified 1179 septin-associated proteins. Interaction candidates were cross-validated using high-throughput yeast two-hybrid (Y2H) screening (Figure 5) and further tested by targeted Y2H and co-expression studies.

To identify proteins associated with septins during infection-related development, we implemented a temporal discovery proteomics workflow based on co-immunoprecipitation-mass spectrometry (Co-IP-MS) using GFP-tagged Sep3, Sep4, Sep5, and Sep6, together with a cytoplasmic GFP control (Figure 1C). Immunoprecipitations were performed from spores, germinated spores, and appressoria across six time points (0, 2, 4, 8, 16, and 24 h) as well as from mycelium. This analysis identified 1179 putative septin-associated proteins, including a high-confidence set of 349 interactors reproducibly detected across three biological replicates (Table S1 and S2). Hierarchical clustering revealed distinct temporal interaction profiles for each septin (Figure S1). Several interactors displayed stage-specific enrichment coinciding with septin polymerization and ring maturation, indicating that the septin interactome undergoes dynamic remodelling during the transition from germ-tube growth to appressorium maturation. Expression and recovery of GFP-tagged septins were confirmed by MS-based quantification (Table S3), validating stable expression and efficient immunoprecipitation and supporting robust comparison across developmental stages.

### Stage-specific recruitment of septin-associated proteins occurs during appressorium development

Quantitative analysis of the Co-IP-MS dataset showed that Sep4 and Sep5 associate with substantially more interactors than Sep3 or Sep6. Their interaction profiles peaked between 2 and 8 h, declined by 16 h, and increased again at 24 h, mirroring the transition from early appressorium morphogenesis to late-stage repolarization (Figure 2A; Table S4 and S5). Analysis of shared- and unique interactors identified proteins associated with individual septins or specific septin combinations. Prominent subsets included Sep4-Sep5 and Sep4-Sep5-Sep6 interactors, alongside 106 pan-septin-associated interactors (Figure 2B). Network visualization highlighted candidate interactors with established roles in appressorium function and pathogenicity, including enzymes involved in melanin biosynthesis required for turgor generation.^30^ Temporal network analysis further showed dynamic changes in interactor subsets during appressorium maturation, indicating extensive remodelling of the septin interactome in parallel with morphogenetic progression (Figure 2D).

To compare infection-specific and vegetative associations, we performed Co-IP-MS and fluorescence localisation analyses in mycelial hyphae. Sep4 and Sep5 localised predominantly to the plasma membrane, including septa and sites of positive curvature, whereas Sep3 and Sep6 displayed greater cytoplasmic distribution, with Sep6 enriched sub-apically near the Spitzenkörper (Figure S2A). During vegetative growth, Sep6 associated with the largest number of interactors, followed by Sep4 and Sep5, while Sep3 showed fewest associations (Figure S2B). Intersection analysis identified both septin-specific interactors and shared subsets (including Sep4-Sep6, Sep4-Sep5-Sep6, and all four septins), consistent with septins engaging developmentally distinct protein networks organised around a shared set of interactors (Figure S2C).Comparative analysis of interactors identified during appressorium development and vegetative growth for each septin revealed condition-specific and shared subsets, including interactors unique to each condition as well as a conserved set present across both developmental stages (Figure S2D).

### Temporal clustering reveals canonical and novel septin functions during appressorium development

To resolve the temporal organisation of septin-associated protein networks, interactors were grouped based on the developmental time point at which the protein reached peak abundance (Figure 3). This analysis revealed that septin interactions are highly stage-specific, with most proteins showing peak association at a single time point, thereby forming temporally distinct interaction modules during appressorium development.

Across all four septins, gene function enrichment analysis identified a conserved set of interactors maintained throughout the time course, linking septins to canonical functions including cytoskeletal organisation, polarity maintenance, and intracellular trafficking (Figure 3). This conserved network included actin (MGG_03982), α/β-tubulins (MGG_06650/MGG_11412), the polarity regulator Cdc42 (MGG_02731), the cell-end marker Tea1 (MGG_02875), and exocyst component Sec6 (MGG_03235), consistent with stable associations with fundamental cellular processes.

**Figure 2.**
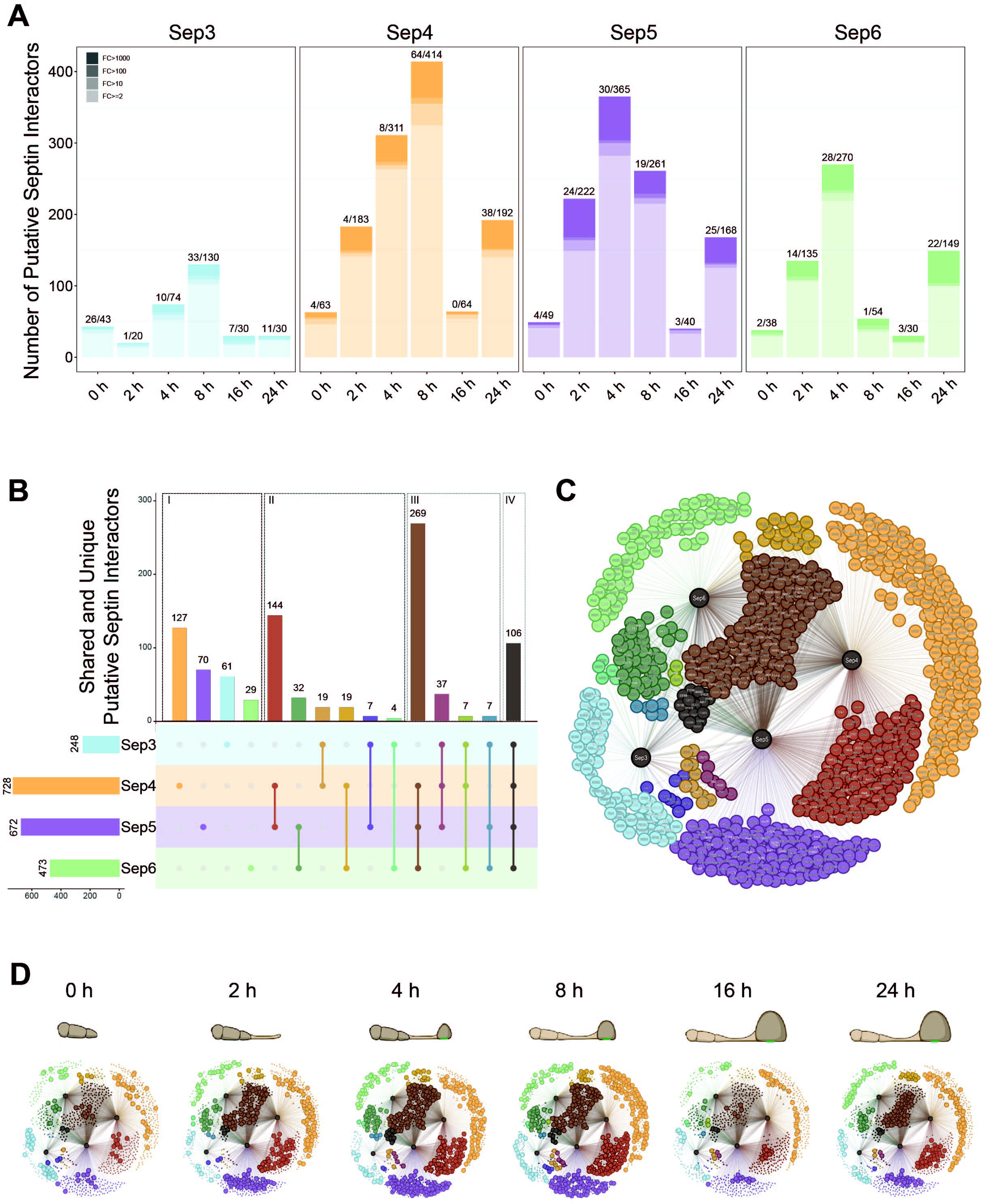
Quantitative and network-based analysis reveals dynamic composition and temporal organisation of the septin interactome during appressorium development. (A) Quantitative summary of putative septin interactors identified for Sep3, Sep4, Sep5, and Sep6 across six developmental time points (0, 2, 4, 8, 16, and 24 h). Bar heights indicate the total number of interactors per septin and time point, with segmented shading representing different fold-change thresholds (FC ≥ 2, > 10, > 100, and > 1000). The number of significant interactors (p < 0.05) and total number of interactors at each time point are indicated above corresponding bars. (B) UpSet plot illustrating shared and unique interactors among the four core septins, organized into four major categories: (I) interactors associated with a single septin, (II) interactors shared by two septins, (III) interactors shared by three septins, and (IV) interactors common to all four septins. Lines connecting septin nodes below the bars represent the specific combination of septins for each group of interactors. The fifteen interaction patterns displayed include Sep3; Sep4; Sep5; Sep6; Sep3-Sep4; Sep3-Sep5; Sep3-Sep6; Sep4-Sep5; Sep4-Sep6; Sep5-Sep6; Sep3-Sep4-Sep5; Sep3-Sep4-Sep6; Sep4-Sep5-Sep6; Sep3-Sep5-Sep6; and Sep3-Sep4-Sep5-Sep6, each distinguished by a unique colour. (C) Network representation of the *M. oryzae* septin interactome constructed from data in panel B. Each node corresponds to an identified interactor, color-coded according to its specific septin-binding category and annotated with gene names where available. For genes lacking standard names, truncated identifiers derived from MGG locus (e.g., MGG_12345 → 12345) are shown. Significant interactors labelled in white (p < 0.05), and non-significant interactors are shown in grey (p > 0.05). (D) Network analysis illustrating temporal distribution of septin interactors across six pre-penetration developmental stages (0, 2, 4, 8, 16, and 24 h) of *M. oryzae*. As in panel C, each node represents an individual interactor color-coded according to specific septin-binding category. Coloured subnetworks depict dynamic shifts in interactor composition over time, with schematic cartoons above each network representing corresponding stage of appressorium development.

**Figure 3.**
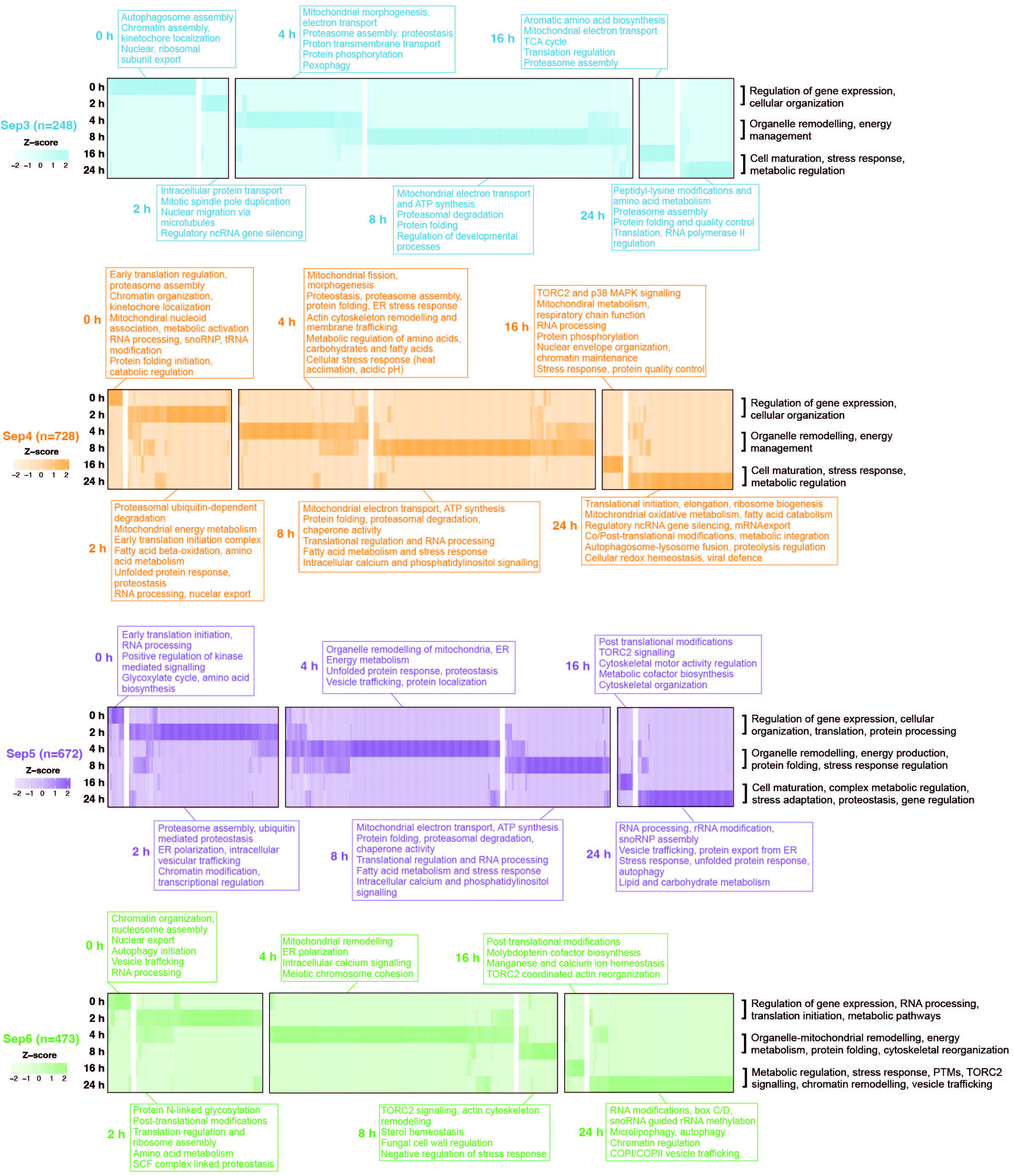
Temporal clustering of septin-associated interactors highlights stage-specific organisation of the interactome during appressorium development. Heat maps of proteins identified as interactors of each core septin - Sep3 (blue), Sep4 (orange), Sep5 (purple), and Sep6 (green) - across six developmental time points (0, 2, 4, 8, 16, and 24 h). Each heat map comprises six rows corresponding to an individual time point, with interactors grouped into six clusters based on peak association. Clusters corresponding to early (0-2 h), mid (4-8 h), and late (16-24 h) developmental stages outlined in black to highlight functional continuity over time. Fold-changes normalised as z-scores (range −2 to 2) for visualisation. Functional annotations indicated in black summarise biological processes enriched within specific temporal windows. Coloured boxes positioned above and below the heat maps highlight time point-specific and previously uncharacterised septin-associated functions within the Sep3, Sep4, Sep5, and Sep6 interactomes.

Stratification into early (0-2 h), mid (4-8 h), and late (16-24 h) stages revealed a clear temporal progression in septin-associated functions. Early-stage modules were enriched for gene expression and cellular organisation, showing septins associating with polarity and nuclear regulatory factors including Bem3 (MGG_09275), a regulator of Cdc42 polarity^31^ identified in genetic interaction screens with septins in yeast^32^, and the nuclear export factor Crm1 (MGG_02526)^33^, also identified in genetic interaction screens with septins.^34^ Additional associations included histone Hho1 (MGG_12797) and ribosome-related components such as Rpp0 (MGG_04467) and the translation factor Tef1 (MGG_03641), supporting links between septins and early transcriptional and translational processes. Mid-stage modules were enriched for organelle remodelling and energy metabolism, spanning ER, mitochondrial, and trafficking systems. These included the ER chaperone Kar2 (MGG_02503), mitochondrial ATP synthase subunits Atp5 (MGG_03152) and Atp4 (MGG_04752), and electron transport components such as Cox5b (MGG_03188) and Cox4 (MGG_01135). Interactions with the AAA-ATPase Cdc48 (MGG_05193), previously identified in genetic interaction screens with septins in yeast,^34^ support conserved functional associations, whereas association with the TCA/redox enzyme Idh1 (MGG_01995) points to septin engagement with mitochondrial metabolism. Septins were also associated with vesicle trafficking and proteostasis factors including Sar1 (MGG_06362), Erv46 (MGG_01245), and Vph1 (MGG_03947), as well as the chaperone Ydj1 (MGG_04462) and the COPI coatomer Ret2 (MGG_05676), reinforcing links to membrane trafficking and protein quality control. Late-stage modules were enriched with metabolic regulation and stress-related processes, including interactions with Adh3 (MGG_03880), identified in genetic interaction screens with septins in yeast,^34^ Idp1 (MGG_07268), Cat2 (MGG_01721), and Ald5 (MGG_03900). Additional associations included the RNA helicase Dhh1 (MGG_03388), the TCA cycle enzyme Kgd1 (MGG_12767), and Hsp70 chaperone Sse2 (MGG_06065), as well as proteins linked to protein turnover and nuclear regulation, such as Pse1 (MGG_03537) and Gus1 (MGG_05956). Further interactions connected septins to chromatin remodelling and cytoskeletal dynamics, including Rvb2 (MGG_03173), the Arp2/3 subunit Arp3 (MGG_03879), and the nitrogen regulator Ure2 (MGG_09138), highlight expanded septin roles in stress adaptation and cellular remodelling. Beyond recapitulating conserved septin-associated processes, temporal resolution of the interactome therefore uncovered stage-specific functional signatures and previously uncharacterised associations, revealing both canonical and expanded roles for septins during appressorium development (Figure 3).

Gene Ontology (GO) enrichment across the developmental time-course provided a systematic view of these functional expansions, identifying diverse metabolic and regulatory processes associated with the septin interactome (Figure 3; Table S4 and S5). Notably, a subset of *M. oryzae* septin-specific interactors, absent from previously reported yeast and mammalian datasets, linked septins to diverse cellular pathways.^35,36^ Sep3 interactors, for example, showed enrichment for mitochondrial metabolism (MGG_01706; GO:0005739), the unfolded protein response and protein folding (MGG_02503; GO:0006457), translation (MGG_09894; GO:0006412), and carbohydrate metabolism (MGG_00530; GO:0005975). Similarly, Sep4, Sep5, and Sep6 exhibit temporal expansions into nuclear, metabolic, and stress-response pathways, each with a distinct enrichment trajectory. Sep4 interactors, for instance, include components of mitochondrial metabolism (e.g., MGG_02617; GO:0005739), ER stress and protein folding (e.g., MGG_02503; GO:0036498), lipid metabolism (e.g., MGG_13767; GO:0006633), and translation (e.g., MGG_05266; GO:0006412), pointing to new potential septin-dependent roles in respiratory metabolism, ER stress adaptation, fatty acid biosynthesis, and protein synthesis. This is reinforced by Sep5 interactors linked to mitochondrial respiration (e.g., MGG_06030; GO:0005739), ER-associated processes (e.g., MGG_02503; GO:0005783), lipid metabolism (e.g., MGG_10492; GO:0006631), and proteostasis (e.g., MGG_05296; GO:0005829). Sep6 interactors, in turn, are associated with mitochondrial remodelling (e.g., MGG_06030; GO:0005739), translation regulation (e.g., MGG_00341; GO:0006412), COPI/COPII-mediated secretion (e.g., MGG_06362; GO:0016192) and ER-associated vesicle trafficking (e.g., MGG_05188; GO:0005783), consistent with septin regulation of organelle dynamics and polarized secretion. Together, these data reveal septin-specific and temporally dynamic functional specialisation, with distinct septins engaging diverse metabolic and regulatory pathways during appressorium development. Collectively, these data define septins as dynamic organisers of temporally structured protein interaction networks during appressorium development, linking conserved cytoskeletal functions to stage-specific metabolic, regulatory, and organelle-associated processes (Figure 3).

### Septins organise virulence-associated proteins during infection

We observed that septin complexes preferentially recruit proteins implicated in appressorium formation and virulence during infection-related development. Analysis of the MagnaGenes *v*. 1.0 database,^37^ revealed septin interactors strongly enriched for previously reported functions required for *Magnaporthe* infection biology. Among the Co-IP-MS interactors with known mutant phenotypes (∼20% of the dataset; n = 67/349 high-confidence, n = 267/1179 total), 15% of high-confidence and 21% of total septin interactors have previously been reported to affect appressorium development (∼3% and ∼5% of all annotated genes), while 63% of high-confidence and 66% of total septin interactors are required for fungal virulence (∼12% and ∼15% overall; Figure 4A).^37^ The remaining ∼80% lack characterized mutant phenotypes, however, suggesting extensive unrecognized virulence functions. Quantitative enrichment analysis further revealed stage-specific interaction patterns across all septins. Proteins implicated in appressorium development peaked between 2-8 h (maximal for Sep4 at 8 h), coinciding with appressorium maturation. Pigmentation enzymes, for instance, were induced at 4-8 h for Sep4/Sep5/Sep6, reflecting melanisation and penetration, whereas virulence-associated proteins were enriched at 4 h and 24 h, matching the onset of host invasion (Figure 4B; Table S6).

**Figure 4.**
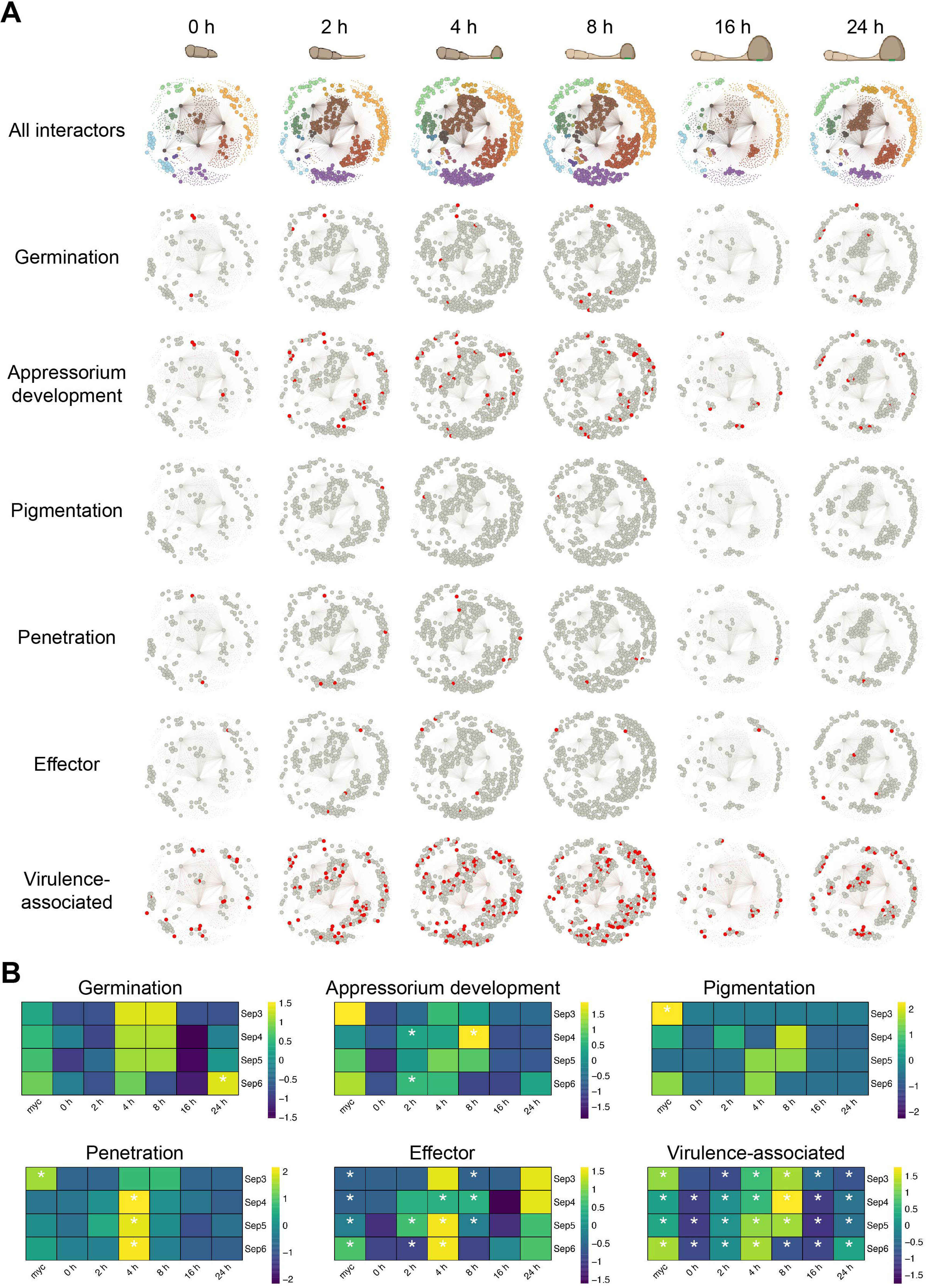
Functional enrichment of septin interactors reveals stage-specific association with virulence-related processes in *M. oryzae*. (A) Temporal network integrating septin interactome data with functional classifications from the publicly available *MagnaGenes* resource. The upper panel depicts time course of appressorium development (0, 2, 4, 8, 16, and 24 h). Below, nine stacked network rows are shown: the top row represents all identified septin interactors (Sep3, Sep4, Sep5, Sep6), color-coded according to septin-interaction combinations as in Figure 2 D, while the subsequent rows display interactors associated with distinct biological processes, including germination, appressorium development, pigmentation, penetration, and effector- and virulence-associated functions. Red-highlighted nodes indicate genes annotated in MagnaGenes as belonging to each process. (B) Quantitative enrichment heat maps showing the relative abundance of interactors associated with six biological processes (germination, appressorium development, pigmentation, penetration, and effector- and virulence-associated functions) for each septin (Sep3, Sep4, Sep5, Sep6) across six developmental time points (0, 2, 4, 8, 16, and 24 h) and during vegetative growth. Colour intensity represents enrichment based on odds ratios, with yellow indicating overrepresentation and dark blue indicating underrepresentation relative to the proteome-wide background. Values are displayed as scaled enrichment scores (range -2 to 2; respectively -1.5 to 1.5), reflecting the degree of over- or underrepresentation compared to *M. oryzae* Guy11 proteome. White asterisks denote statistically significant enrichment (FDR ≤ 0.05, Benjamin-Hochberg correction).

To understand how infection-related septin interactors act mechanistically, we next examined proteins predicted to function as secreted enzymes or annotated as *Magnaporthe* effector proteins (MEPs),^38^ which are implicated in suppression of plant immunity and invasive growth.^39^ This analysis identified a set of high-confidence septin interactors involved in fungal cell wall remodelling and host surface degradation. These include plant cell wall–degrading enzymes such as *MEP364* (MGG_01255), a cellobiose dehydrogenase, as well as fungal cell wall remodelling proteins, including the chitin-modifying enzyme *CBL1* (MGG_09159), GPI-anchored mannoproteins such as *MEP494* (MGG_05575), and oxidative enzymes such as *MEP208* (MGG_05865) and *MEP151* (MGG_10878), which contribute to redox-dependent wall assembly and adhesion.

In parallel, septins associate with components of the ER-Golgi quality control and secretion machinery, including *EMP47* (MGG_05685), *VIP36* (MGG_06367), and protein folding and N-glycosylation factors such as Swp1/OST (MGG_03687), PDIA6 (MGG_06786), glucosidase II (MGG_08623), and N-deacetylase/N-sulfotransferase (NDST; MGG_02390). These factors collectively mediate glycan transfer, protein folding, and quality control, suggesting that septins coordinate secretory cargo maturation prior to delivery.

Additional interactors include chitin synthases *CHS7* (MGG_06064) and *CHS1* (MGG_01802), the O-mannosylation regulator *PTM2* (MGG_07190), and glucan biosynthesis components such as *UGP1* (MGG_01631), linking septins to structural assembly of the fungal cell wall. Together, these data support a model in which septins couple ER-based protein maturation with targeted secretion and localized delivery of cell wall-remodelling enzymes to penetration sites.

### Complementary independent validation of septin interactions during appressorium development

To identify septin interactors using an independent method, we used ultra-high-throughput yeast two-hybrid (Y2H) screening (Hybrigenics). Although we reasoned that Y2H might not identify proteins that interact with higher-order oligomeric septin structures, it does provide a completely separate validation of interactions identified by Co-IP-MS, increasing confidence in their biological relevance. Using this method for Sep3, Sep4, Sep5 and Sep6, we identified 140 putative interactors for the four septins in a cDNA library derived from spores, germinated cells, appressoria and mycelium, revealing a range of direct septin-binding partners (Figure 5A; Table S7). A quantitative comparison showed that Sep5 and Sep4 identified the largest numbers of binary interaction partners (69 and 42, respectively), whereas Sep6 and particularly Sep3 had fewer (25 and 17), mirroring Co-IP-MS appressorium trends and reinforcing the distinct interaction landscapes of each septin (Figure 5B). Most Y2H interactors were septin-specific, consistent with the limitation of the technique in selecting individual protein-protein interactions (Figure 5C). Intersection analysis revealed 1144 proteins uniquely identified by Co-IP-MS, 105 specific to Y2H, and 35 shared candidates, demonstrating the complementarity between native co-complex interactions and binary contacts (Figure 5D; Table S8).

**Figure 5.**
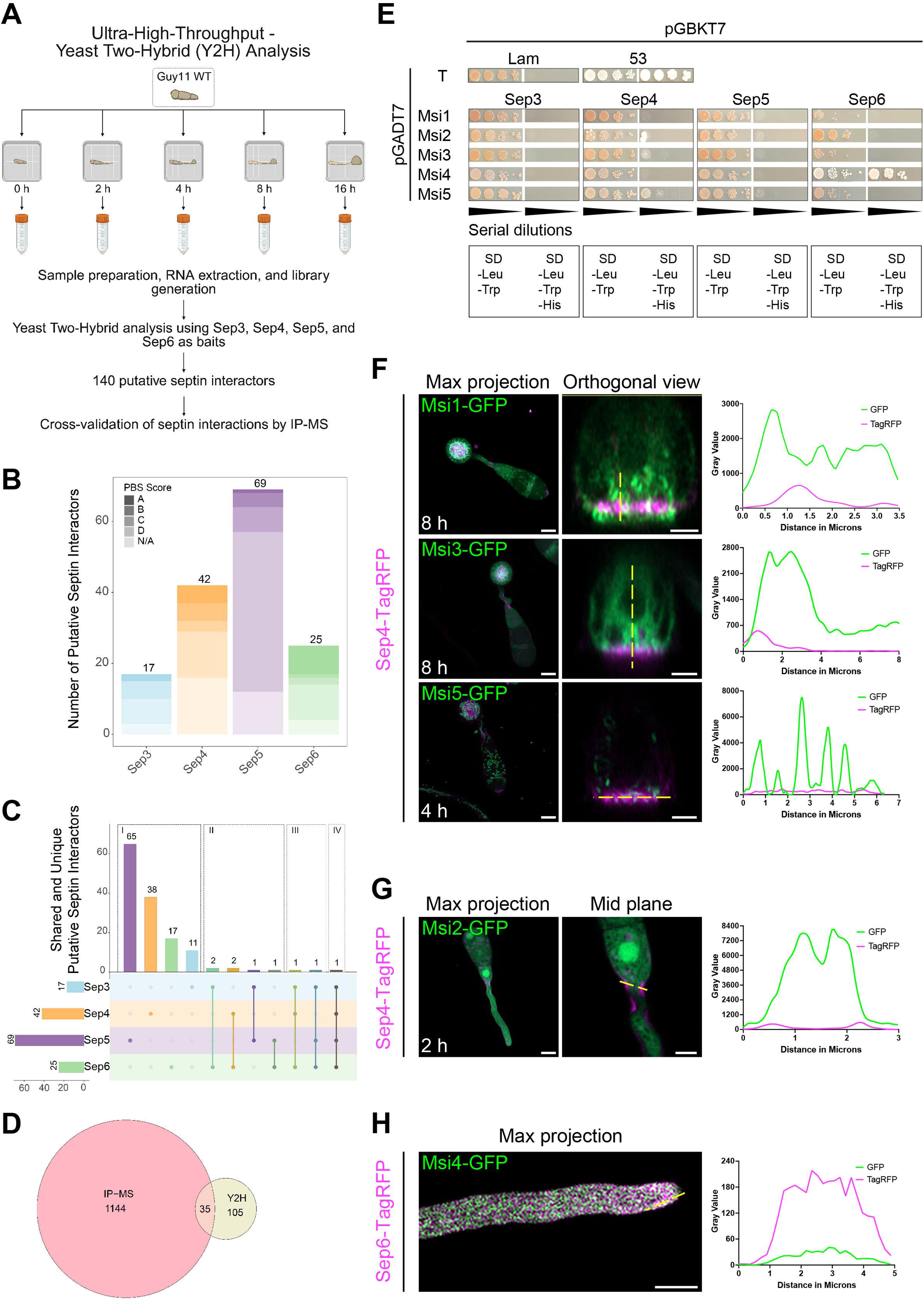
Complementary interaction mapping identifies a conserved set of septin-associated proteins and validates their dynamic association during appressorium development and vegetative growth in *M. oryzae*. (A) Schematic overview of the ultra-high-throughput yeast two-hybrid (Y2H) workflow used to identify septin interactors during appressorium development in *M. oryzae*. A Y2H screen was performed on the wild-type strain Guy11 across multiple developmental stages (0, 2, 4, 8, and 16 h), encompassing conidia, germinated conidia, and appressoria. RNA was extracted from these stages to generate a prey library, which was screened against Sep3, Sep4, Sep5, and Sep6 as bait proteins. The screen was performed in collaboration with Hybrigenics Services and identified 140 putative septin interactors. Candidates were cross-validated against IP-MS dataset (Figure 1C) and subsequently tested by Y2H assays and co-expression analysis. (B) Quantitative summary of putative septin interactors identified for Sep3, Sep4, Sep5, and Sep6. Bar heights represent the total number of interactors detected per septin, with segmented shading indicating different PBS score categories (A, B, C, D, and N/A). The total number of interactors is indicated above each bar. (C) UpSet plot illustrating shared and unique interactors among four core septins (Sep3, Sep4, Sep5, and Sep6). Interactors are categorised into four major groups: (I) unique to a single septin, (II) shared by two septins, (III) shared by three septins, and (IV) common to all four septins. Lines connecting nodes below the bars indicate the specific septin combinations. Eleven distinct interaction patterns are shown (Sep3; Sep4; Sep5; Sep6; Sep3-Sep6; Sep4-Sep6; Sep3-Sep5; Sep5-Sep6; Sep3-Sep4-Sep6; Sep3-Sep5-Sep6; and Sep3-Sep4-Sep5-Sep6) each represented by a unique colour. (D) Venn diagram comparing the septin interactors identified by immunoprecipitation-mass spectrometry (IP-MS) and ultra-high-throughput yeast two-hybrid (Y2H) screening. The overlap highlights high-confidence candidates detected by both merthods and illustrates complementarity between native co-complex capture (IP-MS) and direct binary interaction mapping (Y2H). (E) Targeted Y2H assays validating interactions between five candidate *<u>M</u>agnaporthe* <u>s</u>eptin <u>i</u>nteractors (Msi1- Msi5), identified in both IP-MS and Y2H datasets, and the four core septins (Sep3, Sep4, Sep5, and Sep6). Prey constructs expressing candidate proteins were co-expressed with bait constructs containing individual septin. Interactions were assessed by growth on selective synthetic defined (SD) medium lacking tryptophan, leucine, and histidine (SD -Trp -Leu -His), indicating reporter activity. Control growth on non-selective SD medium lacking tryptophan and leucine (SD -Trp -Leu) confirmed transformation efficiency. Yeast cultures were spotted as 10-fold serial dilutions to assess interaction strength. (F) Confocal laser scanning microscopy combinded with Airyscan super resolution of candidate septin interactors Msi1, Msi3, and Msi5 tagged with GFP and co-expressed with Sep4-TagRFP during appressorium development (8 h). For each protein (left to right), images show: a maximum intensity projection, an orthogonal view illustrating spatial organisation relative to the appressorium base; and corresponding line-scan fluorescence intensity profiles quantifying the spatial distribution of GFP and TagRFP signals. Yellow dotted lines indicate the regions analysed in line scans. Scale bars: 5 μm (maximum intensity projections) and 2 μm (orthogonal views). (G) Confocal laser scanning microscopy (Airyscan super resolution) analysis of Msi2-GFP co-expressed with Sep4-TagRFP during germination (2 h). Images (left to right) show a maximum intensity projection, a mid-plane optical section used for line scan analysis, and the corresponding line scan profile quantifying spatial separation of GFP and TagRFP signals. Yellow dotted lines indicate analysed region. Scale bars: 5 μm. (H) Live cell imaging using confocal laser scanning microscopy of *M. oryza*e expressing Msi4-GFP co-expressed with Sep6-TagRFP during vegetative growth. Images show a maximum intensity projection and a corresponding line scan profile quantifying spatial separation of fluorescence signals. Yellow dotted lines indicate analysed region. Scale bar: 5 μm.

To select the most stringent and confidently assigned septin interactors, we selected five *<u>M</u>agnaporthe* <u>s</u>eptin <u>i</u>nteractors (Msi1-Msi5) from a group of 35 shared candidates for targeted analysis. Full-length individual Y2H assays confirmed specific interactions identified in the ultra-high-throughput screen: Msi1 (MGG_01508) with Sep4; Msi2 (MGG_05155) with Sep4/Sep5; Msi3 (MGG_06316) with Sep4; Msi4 (MGG_10677) with Sep6; and Msi5 (MGG_13781) with Sep4/Sep5, as shown in Figure 5E. Live-cell imaging of GFP-tagged Msi1-Msi5 co-expressed with Sep4-TagRFP (Msi1-Msi3, Msi5) or Sep6-TagRFP (Msi4) showed dynamic co-localisation patterns in their native cellular context, consistent with temporal Co-IP-MS predictions and linked each Msi to distinct infection-related developmental stages (Figure 5F-H, S3). Msi1, a predicted BAR-domain protein homologous to yeast Gvp36,^40^ initially displayed cytoplasmic and punctate localisation in conidia and germinating cells before assembling into a disc-like structure at the appressorium base. This disc, slightly larger than the septin ring, peaked in intensity around 8 h, consistent with previous studies.^41^ Orthogonal imaging and line-scan analyses confirmed that the Msi1 disc is positioned just beneath the cortical septin assembly at the appressorium base, consistent with its temporal interaction profile predicted by Co-IP-MS (Figure 5F, S3A). Msi2, a predicted adenosylhomocysteinase that interacts with septins at early stages, exhibited dynamic localisation matching Co-IP-MS predictions. It primarily formed cytoplasmic round structures from 0-4 h but strikingly colocalised with Sep4 at sites of positive membrane curvature in germ tubes at 2 h. During appressorium development, Msi2 transitioned to filamentous assemblies before reverting to round structures (Figure 5G, S3A). Msi3, predicted to encode a curved DNA-binding protein, exhibited distinct spatial dynamics that mirrored its Co-IP-MS-predicted interaction with septins at 8 and 24 h. In conidia, Msi3 localised as small cytoplasmic puncta, transitioning in germinated cells to a network-like cytoplasmic distribution. Within the appressorium, it formed an intricate network interwoven with the upper surface of the septin disc around 8 h, before progressively condensing into a distinct disc at the appressorium base between 16 and 24 h (Figure 5F, S3A). Msi4, a predicted mannose-6-phosphate isomerase that interacts with Sep6 during vegetative growth, predominantly localised in the cytoplasm of growing hyphae but strikingly colocalised with Sep6 at hyphal tips, a site critical for polarized growth (Figure 5H, S3B). Line-scan analyses revealed overlapping fluorescence profiles despite stronger Sep6 signal intensity, suggesting functional coupling of Msi4 with septin assemblies at growth-associated sites. Msi5, a putative serine hydroxymethyltransferase predicted to interact with septins at 4, 8, and 24 h, exhibited progressive structural organisation during appressorium development. In conidia, Msi5 displayed punctate and tubular fluorescence, forming elongated filamentous structures during germination and early germ tube growth. At 4 h, Msi5 transitioned to punctate foci, and during appressorium maturation formed increasingly filamentous structures aligned with the growth axis inside germ tubes. Orthogonal imaging and line-scan analyses confirmed partial incorporation of Msi5 into the septin disc at 8 h, suggesting structural integration with septin scaffolds during appressorium morphogenesis (Figure 5F, S3A). When considered together, the spatial and temporal co-localisation of these metabolic enzymes, BAR-domain proteins, and DNA-binding proteins with septin assemblies highlights septins as dynamic platforms that coordinate metabolic processes, cytoskeletal architecture, and membrane remodelling in *M. oryzae*.

### Functional characterization of the septin interactor Msi1 reveals a role in fungal invasive growth

To assess the biological significance of septin interactors in fungal pathogenicity, we focused on Msi1, a predicted BAR-domain protein homologous to the yeast membrane curvature sensor Gvp36.^40^ Live-cell imaging revealed that Msi1 dynamically re-localizes during infection-related development, initially displaying cytoplasmic and punctate fluorescence in conidia and germinating cells before assembling into a distinct disc-like structure at the base of the appressorium. This Msi1 disc, slightly larger than the septin ring, peaked in intensity around 8 h after spore germination. Orthogonal views and line-scan analyses confirmed that Msi1 is positioned immediately beneath the cortical septin assembly, consistent with its temporal interaction profile identified by Co-IP-MS (Figure 5F, S3A, 6A). Super-resolution microscopy further resolved the Msi1 disc as a mesh-like network with prominent fluorescent puncta incorporated into the structure and stronger signal intensity concentrated at its periphery (Figure 6B).

**Figure 6.**
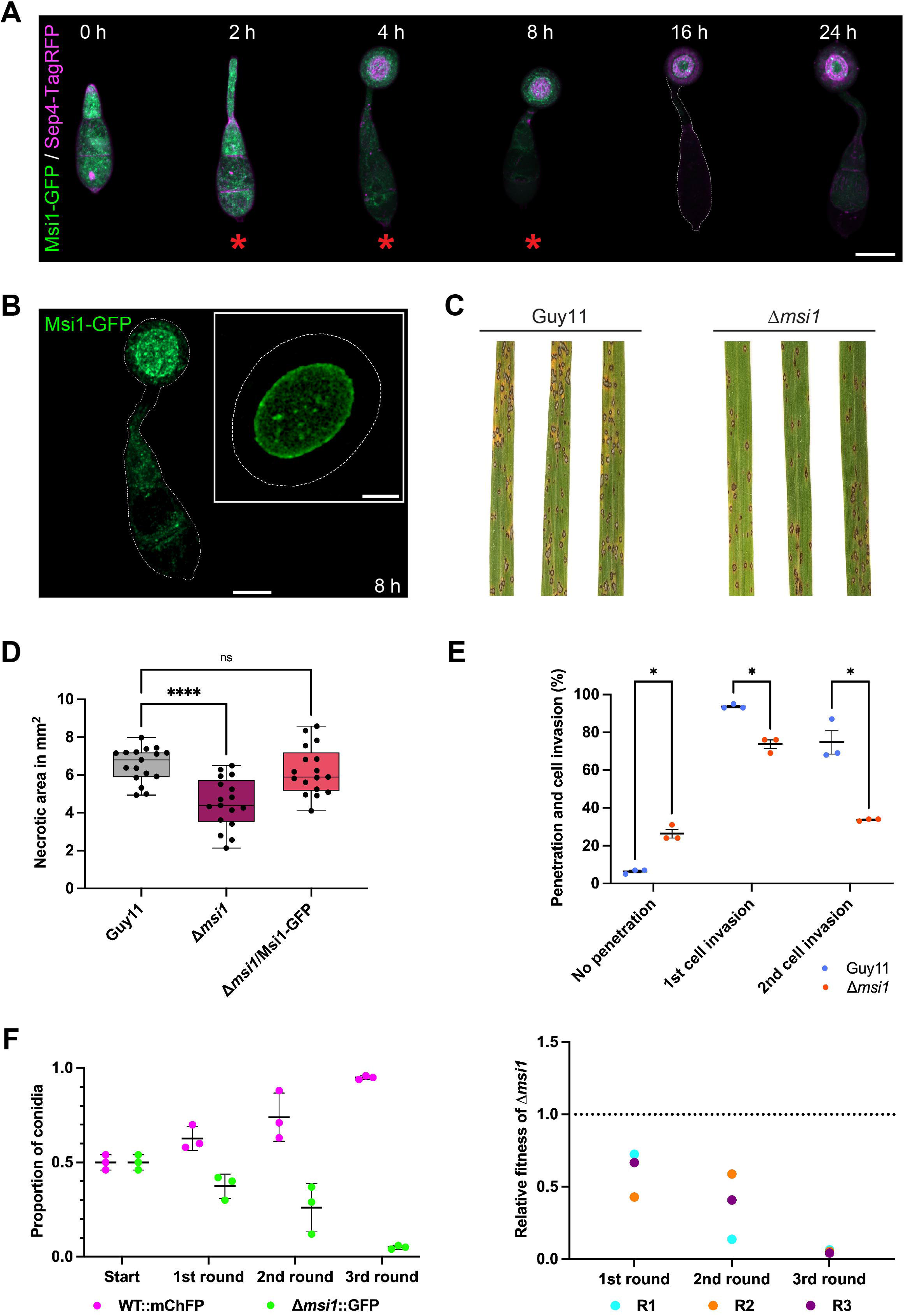
The septin interactor Msi1 localises to the appressorium pore and is required for virulence of *M. oryzae*. (A) Confocal laser scanning microscopy combined with Airyscan super-resolution imaging shows the dynamic localisation of Msi1-GFP co-expressed with Sep4-TagRFP during appressorium development (0, 2, 4, 8, 16, and 24 h post inoculation). Red asterisks indicate time points at which interaction with Sep4 was detected in IP-MS dataset. (B) Confocal laser scanning microscopy combined with Airyscan super-resolution imaging shows localisation of Msi1-GFP as a disc-like structure at the base of the appressorium at 8 h post inoculation. An inset Structured Illumination Microscopy (SIM) image provides enhanced super-resolution visualisation of the Msi1-GFP disc. Scale bars: 5 μm (main image) and 2 μm (inset). (C) Rice seedlings (cultivar CO-39) were spray-inoculated with equal concentrations (1 × 10^5^ conidia/mL) of Guy11 (left) and Δ*msi1* (right) strains and incubated for 5 days. Representative infected leaves are shown. (D) Quantification of necrotic lesion area (mm^2^) from leaf drop assays using 3-week-old rice seedlings (cultivar CO-39) inoculated with equal concentrations (1 × 10^5^ conidia/mL in 0.2% Gelatin) of Guy11 WT, Δ*msi1*, and complemented Msi1-GFP strains. Seedlings were incubated at 26°C for 5 days. Box plots show lesion area (mean ± SE) from a representative experiment. Infections were performed independently three times with similar results. Asterisks indicate statistically significant differences compared to WT (ordinary one-way ANOVA, p ≤ 0.05). (E) Quantification of penetration and host cell invasion rates (%) from leaf sheath assays using 3-week-old rice seedlings (cultivar CO-39) inoculated with equal concentrations (1 × 10^3^ conidia/mL in 0.2% Gelatin) of Guy11 WT, Δ*msi1*, and complemented Msi1-GFP strains. Seedlings were incubated at 26°C for ∼48 h. Infection progression was assessed by brightfield microscopy. Violin plots show penetration and invasion rates (mean ± SE) from a representative experiment. Host invasion assays were independently repeated three times with similar results. Asterisks indicate statistically significant differences compared to WT (Multiple unpaired t tests, p ≤ 0.05). (F) Relative pathogenic fitness assay measuring competitive growth of Guy11 expressing mCherry (WT::mChFP) and Δ*msi1* expressing GFP (Δ*msi1*::GFP) over three serial generations on barley (cultivar Golden Promise). Dot plot shows strain proportions recovered per generation (left), and relative fitness coefficient values of Δ*msi1*::GFP in each generation (right). Data represent mean ± SE from a representative experiment. Experiments were independently repeated three times with similar results.

Phylogenetic analysis of fungal BAR-domain proteins revealed Msi1 to belong to a conserved clade present in species exhibiting diverse fungal lifestyles, including biotrophs, hemibiotrophs, necrotrophs, endophytes, and saprophytes, with seven of these BAR domain proteins recovered in our septin interactome dataset (Figure S4A). To test the function of Msi1, we generated a targeted deletion mutant of *M. oryzae* Δ*msi1* which was confirmed by whole-genome sequence analysis (Figure S4B and C). Phenotypic characterization revealed that Δ*msi1* mutants germinated normally and showed no major growth defect on minimal medium but exhibited a significant reduction in growth at varying temperatures, indicating a conditional growth defect (Figure S4D-G).

Functional assays demonstrated that deletion of *MSI1* significantly impairs virulence (Figure 6). Spray inoculation of rice seedlings and droplet inoculation of barley leaves with Δ*msi1* mutants produced markedly fewer and smaller necrotic disease lesions than the isogenic wild-type Guy11 strain (p ≤ 0.05), and reintroduction of *MSI1* restored wild-type pathogenicity (Figures 6C, 6D). Leaf-sheath assays confirmed these observations, revealing reduced penetration and limited progression beyond the first host cell by Δ*msi1* appressoria relative to Guy11 (Figure 6E). To further quantify the virulence defect, we performed a competitive fitness assay^38^ in which *Δmsi1* mutants expressing cytosolic GFP were co-inoculated with an isogenic wild-type strain expressing cytosolic mCherry onto rice seedlings. Mixed infections were recovered and re-inoculated across three successive plant infection cycles to simulate serial host colonization. Fluorescence quantification showed a progressive decline in the GFP:mCherry signal ratio, demonstrating that Δ*msi1* mutants were rapidly outcompeted by the wild type, declining to ∼5% of the wild-type population after three successive infection cycles, indicating a severe loss of relative fitness during host infection (Figure 6F).^38^

Notably, Msi1 localises to mobile punctate structures not only at the appressorium base but also within invasive hyphae during plant infection, as shown by live-cell imaging and supported by video microscopy (Figure S3B, S4H; Video 1). This dynamic behaviour is consistent with a role for Msi1 in membrane remodelling or signalling processes critical during host colonization.

Collectively, these findings validate the utility of the time-resolved septin interactome generated in this study to identify new septin-dependent gene functions. Msi1 is a key septin-associated protein that organises a specialized membrane-associated disc at the appressorium base and is necessary for effective host penetration in *M. oryzae*. Given the broad conservation of BAR-domain proteins and their enrichment among septin-associated virulence factors (Figure 4), Msi1 likely represents a link between septin-mediated cytoskeletal organisation, membrane-curvature sensing, and infection-structure differentiation.

## DISCUSSION

Fungal pathogens pose a major threat to global health and food security, infecting plants, insects, and humans through specialised infection structures and coordinated virulence programmes. These processes require tight integration of cytoskeletal organisation, membrane remodelling, secretion, and metabolic adaptation to enable host invasion and colonisation.^3,27,42^ Among conserved regulators of cellular architecture, septins are key cortical GTP-binding proteins that organise polarity and cytokinesis. Septins are evolutionarily conserved across opisthokonts.^3,5,10^ but play central roles in infection-related morphogenesis in fungi. Septins are essential for host penetration in fungal pathogens, most prominently in the rice blast fungus *Magnaporthe oryzae*, one of the most devastating plant pathogens worldwide, where they assemble into a ring at the appressorium pore that enables mechanical rupture of the host surface during pathogenesis.^24,27^

Despite this central role, our understanding of how septins execute their biological functions remains limited, particularly at the level of their dynamic protein interaction networks. While previous work has defined septin localisation and higher-order assembly, the composition, temporal remodelling, and functional organisation of septin-associated proteins during infection-related development remain largely unresolved. As a result, it is unclear how septins integrate cytoskeletal architecture with organelle dynamics, metabolic pathways, and signalling processes to coordinate morphogenesis and virulence at a systems level. Addressing this gap is essential to understand how septins function as dynamic organisers of infection-related development and may reveal new targets for controlling fungal disease.

In this study, we combined time-resolved co-immunoprecipitation mass spectrometry with high-throughput yeast two-hybrid screening to define a stage-resolved septin interactome during appressorium development. Our analysis reveals extensive temporal rewiring of septin-associated networks, identifying a conserved septin interactome alongside dynamic modules that link septins to metabolism, organelle function, secretion, and stress adaptation.

The first major conclusion of this study is that septin organisation during infection-related development is highly dynamic and accompanied by extensive remodelling of the septin interactome. Super-resolution imaging revealed that the four core septins of *M. oryzae* undergo a progressive transition from dispersed, filamentous and punctate structures during early germination to a disc-like assembly and ultimately a mature basal ring at the appressorium pore, reinforcing the idea that higher-order septin organisation provides mechanical and spatial control of appressorium-mediated infection. This structural progression is mirrored by a temporally resolved interactome comprising 1179 putative septin-associated proteins with a high-confidence set of 349 interactors, which display pronounced stage-specific recruitment during appressorium development. Hierarchical clustering and quantitative analyses revealed distinct temporal association profiles for individual septins, with all four core septins displaying a shared peak in interactor recruitment between 2 and 8 h post-inoculation, followed by a decline at 16 h and subsequent reconfiguration at 24 h, while Sep4 and Sep5 consistently recruited a greater number of interactors overall. In addition to temporal rewiring, the septin interactome is organised into shared and septin-specific subsets, comprising a conserved set of pan-septin interactors alongside proteins associated with individual septins and subsets of defined multi-septin assemblies. This organisation is consistent with the hetero-oligomeric architecture of septin complexes, in which distinct septin subunits contribute both shared and subunit-specific interaction interfaces, thereby enabling selective recruitment of proteins to particular septin assemblies. Comparison of infection-related development with vegetative growth further revealed that septin interactomes comprise both shared and condition-specific components, including a conserved set of interactors present during both vegetative growth and infection-related development, alongside distinct subsets unique to each developmental state. Together, these findings challenge the prevailing view of septins as static scaffolds or diffusion barriers and instead support a model in which septins act as dynamic, systems-level organisers that coordinate distinct protein-protein interaction networks in a stage-dependent manner during morphogenesis.^3^

The second major conclusion of this study is that septins coordinate a broader and more dynamically organised range of cellular processes during infection-related development than previously appreciated. Temporal clustering reveals that each core septin assembles a strongly stage-resolved interactome, with many interactors confined to distinct developmental time points during appressorium morphogenesis, while a persistent set links all septins to canonical functions in cytoskeletal organisation, polarity maintenance, and vesicle trafficking. Notably, this conserved set includes well-established septin-associated components such as actin and tubulins, as well as polarity regulators including Cdc42 and the polarity landmark protein Tea1, consistent with established roles of septins in cytoskeletal organisation and polarity control.^43,44^ Beyond this expected set, the septin interactome encompasses a broad range of additional proteins and cellular functions, including factors associated with secretion and deployment of virulence determinants during plant infection. This expansion also includes several *M. oryzae*-specific interactors with no clear homologues in yeast or mammalian septin datasets, pointing to lineage-specific adaptations linked to infection-related development.

Septin-associated networks follow a strikingly ordered developmental trajectory that mirrors appressorium differentiation. Early-stage septin interactors include regulators of polarity, gene expression, and chromatin organisation, such as the Cdc42 regulator Bem3 and the nuclear export factor Crm1, suggesting a role in coordinating initial cellular reprogramming. Mid-stage septin networks expand to include organelle remodelling, membrane trafficking, and energy production, exemplified by interactions with mitochondrial ATP synthase components and ER chaperones such as Kar2, consistent with the increasing bioenergetic and biosynthetic demands of appressorium maturation. Late-stage interactors are enriched for metabolic enzymes and stress-associated factors, including aldehyde dehydrogenases and RNA-associated proteins, reflecting the transition to a mature, stress-adapted infection structure primed for host invasion. This temporal progression, from early organisation to bioenergetic and organelle restructuring and finally to stress adaptation, recapitulates functional transitions required to build and activate a melanised, turgor-generating appressorium that facilitates host penetration.^27,45–47^

Strikingly, many of the processes captured within the septin interactome extend beyond their traditionally described roles in cytoskeletal organisation and polarity, revealing connections to mitochondrial metabolism, lipid and fatty acid metabolism that underpin turgor generation in the appressorium, ER homeostasis and the unfolded protein response, and vesicle-mediated secretion, including COPI- and COPII-dependent trafficking required for polarised growth and host penetration. Additional associations include protein-folding and proteostasis, nuclear export, RNA regulation, and chromatin organisation, further highlighting the breadth of septin-associated functions. Together, these data define a high-confidence set of septin-associated proteins while also revealing a broader functional landscape that points to additional layers of septin-associated biology. While some associations arise from lower statistical confidence, they remain biologically coherent and highlight processes such as cellular redox homeostasis, amino acid biosynthesis, protein phosphorylation, fungal cell wall regulation, and more specialised pathways including regulatory ncRNA networks and kinetochore-associated functions, many of which are consistent with the extensive metabolic and cellular reprogramming required during appressorium development. In this context, these extended interactions suggest that septins contribute to integrating the physiological and structural transitions underlying infection. Together, these findings reveal how a broad spectrum of cellular pathways, including several previously underappreciated in septin biology, are integrated and deployed in a temporally structured manner, supporting a model in which septins act as dynamic organisers that coordinate structural, metabolic, and regulatory processes during infection-structure differentiation.

An independent line of evidence further supports the robustness and biological relevance of the septin interaction landscape defined here. Ultra-high-throughput yeast two-hybrid screening identified a complementary set of septin-binding proteins across multiple developmental stages, extending the Co-IP-MS dataset by capturing direct binary interactions that may be underrepresented in native co-complexes. Despite limited numerical overlap between the two approaches, the shared subset of interactors defines a high-confidence candidates and underscores the complementarity of these methods to resolve both stable and transient septin associations.

From this intersecting dataset, five *M. oryzae* septin interactors (Msi1-Msi5) were selected for targeted validation. Directed interaction assays confirmed specific associations with individual core septins, while live-cell imaging demonstrated that these proteins co-localise with septin assemblies in a spatially and temporally dynamic manner during appressorium development and vegetative growth. These interactors span diverse functional classes, including metabolic enzymes and a predicted membrane-curvature-associated BAR-domain protein, reflecting the breadth of processes identified in the global dataset. Notably, their localisation patterns closely mirrored temporal interaction profiles predicted by Co-IP-MS, providing strong experimental support for stage-resolved organisation of septin-associated networks. For example, Msi1 assembled into a disc-like structure positioned cortical to the septin ring at the appressorium base, supporting a role in membrane remodelling at sites of positive curvature. Msi5 showed partial incorporation into septin assemblies during key morphogenetic stages, linking metabolic activity to structural organisation, while Msi4 co-localised with septins at hyphal tips, supporting a role in polarised growth during vegetative development. Together, these observations demonstrate that septin-associated proteins are not only physically linked to septin assemblies but dynamically integrated into distinct cellular structures in a developmentally-regulated manner.

Collectively, these validation experiments provide direct evidence that the septin interactome defined here reflects biologically meaningful and spatially organised interaction networks. By linking septins to proteins involved in metabolism and membrane dynamics, these findings reinforce a model in which septins act as dynamic organisers that coordinate diverse cellular processes across both infection-related development and vegetative growth in *M. oryzae*.

A key implication of the septin interaction landscape defined here is its strong functional coupling to infection biology and virulence. Integration with the MagnaGenes *v*.1.0 database^37^ revealed that septin-associated proteins are strongly enriched for factors required for appressorium development and pathogenicity relative to their representation in the genome. Notably, 63-66% of septin interactors are associated with virulence phenotypes, compared to ∼12-15% genome-wide. Similarly, 15-21% of interactors affect appressorium development, relative to ∼3-5% of annotated genes genome-wide. A substantial proportion of interactors therefore influence virulence outcomes, while the majority remain functionally uncharacterised, pointing to an extensive, previously unrecognised virulence network. Temporal analysis further demonstrated that these interactions are dynamically deployed during infection-related development, with distinct cohorts enriched at stages corresponding to appressorium formation, melanisation, and host invasion.

Mechanistically, these infection-associated interactors define a coordinated network that links septin organisation to cell surface remodelling, secretion, and membrane dynamics required for appressorium function. In particular, septins associate with proteins involved in fungal cell wall remodelling, host surface degradation, and ER-Golgi-mediated processing and delivery, collectively supporting the assembly of a specialised secretion system necessary for host penetration. This includes enzymes mediating chitin and glucan synthesis, regulators of cell wall architecture, and quality-control machinery that ensures proper folding, modification, and targeting of secreted proteins. Such septin-dependent spatiotemporal coordination is likely essential for generating the mechanically robust and biochemically active cell surface required for turgor-driven penetration and host invasion.

The biological significance of this organisation is underscored by functional analysis of the septin interactor Msi1. This BAR-domain protein forms a distinct membrane-associated structure at the appressorium base and displays dynamic localisation during both development and invasive growth. Loss of Msi1 results in a pronounced reduction in virulence, including defects in host penetration, invasive growth, and competitive fitness, directly linking septin-associated membrane organisation to pathogenic success. However, the relatively moderate virulence phenotype of the Δ*msi1* mutant, despite clear defects in infection, is consistent with the presence of multiple BAR domain proteins in the genome, including seven identified within the septin interactome, which likely confer partial functional redundancy. Msi1 therefore likely exemplifies a broader group of septin-associated BAR domain proteins that link septin organisation to membrane remodelling during infection.

When considered together, these findings establish septins as central organisers of infection-related processes, coordinating spatial and temporal deployment of proteins required for appressorium function and host invasion, and identify the septin interaction network as a critical determinant of fungal pathogenicity, consistent with the non-pathogenic phenotype of septin mutants.^24^ However, our findings support a revised conceptual framework for septin function during fungal infection. Rather than acting solely as static scaffolds or diffusion barriers, septins emerge as dynamic organisers that integrate cytoskeletal architecture with membrane remodelling, secretion, and metabolic regulation to coordinate complex cellular state transitions. By assembling stage-specific interaction networks, septins provide a spatiotemporal platform that links structural organisation with the biochemical and physiological processes required for morphogenesis and host invasion. This systems-level view of septin function is likely to extend beyond fungal pathogenesis and may reflect a more general principle of septin biology across fungi and animals. The dynamic interaction landscapes defined here further indicate that septins organise context-dependent protein networks rewired in response to developmental and environmental cues, positioning them as integrative organisers that link cellular architecture with cellular physiology.

Importantly, the comprehensive, temporally resolved septin interactome presented here provides a resource for the wider septin research community and a framework for identifying conserved and lineage-specific components of septin-associated biology across fungi and animals. The large number of previously uncharacterised septin-associated proteins identified in this study, for example, many of which are linked to virulence, provides a foundation for uncovering new septin-dependent mechanisms.

## Limitations of the study

While this study provides a comprehensive, temporally resolved view of septin-associated protein networks during infection-related development, there are limitations to the study. First, our analyses were performed using in vitro-induced appressoria on artificial hydrophobic surfaces, which, although well established for studying infection-related morphogenesis, may not fully capture the complexity of host-derived signals encountered during plant infection. Investigating septin interactions directly in infected plant tissue would provide valuable additional insight but remains technically challenging due to limited fungal biomass and interference from host-derived proteins, and our attempts to achieve sufficient depth in Co-IP-MS experiments under these conditions have not yet been successful. Second, as with all proteomic interaction studies, the dataset likely contains a proportion of false-positive or indirect associations arising from co-complex purification. We attempted to mitigate this through stringent quality control, replicate consistency, quantitative filtering, and orthogonal validation using yeast two-hybrid screening, to support the robustness of the identified interaction landscape. The complementary nature of the approaches used here also highlights that different methods capture distinct classes of interactions, including both stable assemblies and transient or direct binary associations. We know that focusing on septin interactors identified by Co-IP-MS and Y2H analysis excluded many important interactors of the assembled septin hetero-polymers present in appressoria, but in this initial study we chose to focus solely on such interactors because they have highest possible confidence scores. We do intend in future, however, to adopt a wider functional analysis of septin interactors that focuses much more on Co-IP-MS data sets, which likely provide the most valuable resource. Finally, functional validation was necessarily limited to only a small subset of septin interactors, and further studies will be required to define mechanistic roles of the large number of uncharacterised septin-associated proteins identified.

## ACKNOWLEDGEMENTS

We thank Dr. Eva Wegel (JIC) for providing technical support on microscopy. This work was supported by grants to N.J.T. from the European Research Council Advanced Grant SEPBLAST awarded under the UKRI guarantee scheme (EP/X022439/1), the Gatsby Charitable Foundation, and The Biotechnology and Biological Sciences Research Council Institute Strategic Project in Advancing Plant Health (BB/Y002997/1).

## AUTHOR CONTRIBUTIONS

N.J.T. and I.E. conceived the project. I.E., N.S., M.G.R., P.D., F.L.H.M. and W.M. performed experiments and analysed the data. I.E., N.S. and M.G.R. prepared figures and tables. N.S. conceived, designed and performed computational analyses and developed the code repository. I.E., M.G.R. and N.J.T. wrote the manuscript, with contributions from all authors.

## METHODS

### KEY RESOURCES TABLE

see Table S10

### RESOURCE AVAILABILITY

#### Lead contact

Nicholas J. Talbot is the lead contact for interactome-related data, proteomics resources, biological materials and fungal strains.

#### Material availability

All *M. oryzae* strains and plasmids generated in this study are available from the lead contact upon request.

#### Data and code availability

- All proteomics data processing, quality control, visualization, and statistical analyses were performed in R (version 4.3.1, R Core Team 2023).
- Analyses mainly used tidyverse (v2.0.0) for data manipulation, ggplot2 (v3.4.4) for graphics, and custom R scripts for fold change calculations, filtering, and network preparation.
- Processed datasets (LFQ intensities, fold changes, replicate metadata, functional annotations for all four septins across all developmental time points) are provided as tab separated or text files in the Github repository for downstream network and enrichment analyses.
- All custom R scripts used in this study are available at: https://github.com/nehasahu486/Septin-Interactome-in-Magnaporthe-oryzae.
- Any additional information required to reanalyse the data reported in this paper is available from the lead contact upon request.

### EXPERIMENTAL MODEL AND STUDY PARTICIPANT DETAILS

#### *Magnaporthe oryzae* and growth conditions

*Magnaporthe oryzae* strains (wild-type isolate Guy11);^48^ were routinely cultivated on complete medium (CM) agar plates (1.5%) at 25 °C under a 12 h light/dark cycle (Talbot et al., 1993). For long-term storage, strains were grown over sterile filter paper squares (Whatman International) overlaid on CM plates; after growth, paper squares were desiccated and stored at −20 °C.

#### *Oryza sativa* and growth conditions

The blast-susceptible rice (*Oryza sativa*) cultivar CO-39 was used for leaf spray inoculation and leaf sheath assays.^49^ Plants were grown in controlled-environment chambers at 26 °C (day) and 24 °C (night), 16 h photoperiod, and 85% relative humidity.

#### *Hordeum vulgare* and growth conditions

The blast-susceptible barley (*Hordeum vulgare*) cultivar Golden Promise was used for detached leaf and fitness assays.^50^ Plants were grown in a controlled environment cabinet (Sanyo) at 18 °C (day) and 8 °C (night) with a 16 h photoperiod.

### METHOD DETAILS

#### Preparation of conidial samples for *in vitro* appressorium assays, virulence assays and microscopy

##### Conidial harvest for microscopy

Conidia were harvested from 8-12-day-old CM agar plate cultures using a sterile disposable plastic spreader and 1 mL sterile distilled water. Suspensions were filtered through sterile Miracloth (Millipore) and pelleted by centrifugation at 3,500 × g for 5 min at room temperature (Beckman JA-17 rotor). Pellets were resuspended in sterile distilled water, quantified using an Improved Neubauer haemocytometer, and diluted to 5 × 10⁴ conidia/ml for microscopy.

##### Conidial harvest for virulence assays

Conidia were harvested from 8-12-day-old CM agar plate cultures using sterile disposable plastic spreaders and 1 ml sterile distilled water. Suspensions were filtered through sterile Miracloth (Millipore) and pelleted by centrifugation at 3,500 × g for 5 min at room temperature (Beckman JA-17 rotor). Rice leaf sheath assay: Pellets resuspended in 0.2% (w/v) gelatin to 1 × 10³ conidia/ml. Rice leaf spray assay: Pellets resuspended in 0.2% (w/v) gelatin to 1 × 10⁵ conidia/ml. Barley droplet assay: Pellets resuspended in sterile distilled water to 1 × 10⁴ conidia/ml. Fitness assays: Pellets resuspended in sterile distilled water to 1 × 10⁵ conidia/ml.

##### Large-scale conidial harvest for IP-MS and Y2H analysis

Conidia for proteomics and ultra-high-throughput yeast two-hybrid (Hybrigenics) were harvested from 10-day-old CM plates. The fungal material from each CM plate was completely removed using a round cookie cutter (Roys of Wroxham), transferred to a 1 L beaker (≤½ capacity), and just covered with sterile distilled water. Plugs were thoroughly mashed using a sterile potato masher (Roys of Wroxham), poured into a 1 L plastic bottle, and shaken vigorously. The mashed fungal suspension was filtered through sterile Miracloth (Millipore) in a sieve (Roys of Wroxham) over an Erlenmeyer beaker, with gentle squeezing to maximize conidial recovery. The filtrate was passed through sterile Miracloth (Millipore) twice more into 1 L Duran bottles, rinsed to ∼900 mL with sterile water, distributed into 250 ml Falcon tubes, and centrifuged at 1,800 × g (3,500 rpm) for 10 min at room temperature. Supernatants were discarded and pellets resuspended to 7.5 × 10⁵ conidia/ml in sterile water containing 50 ng/ml 1,16-hexadecanediol (Sigma-Aldrich).

#### Confocal and super-resolution microscopy

For confocal laser scanning microscopy (LSCM), conidia of relevant *M. oryzae* strains were incubated on VWR borosilicate cover glasses (22 x 50 mm, thickness no. 1.5) and mounted on VWR microscope slides (VWR International) using Grace Bio-Labs SecureSeal™ imaging spacers (13 mm x 0.12 mm). Imaging was performed at specified time points (0, 2, 4, 8, 16, and 24 hpi) using either a Zeiss LSM 880 Airyscan or a Zeiss LSM 980 Airyscan upright confocal microscope. For the LSM 880, z-stacks were acquired utilizing a multiline argon laser (488 nm for eGFP) and an Airyscan detector. For the LSM 980, micrographs and time-lapse movies were collected using solid-state diode (488 nm for eGFP) and DPSS (561 nm for TagRFP) lasers with an Airyscan 2 detector. Both systems operated with 100x alpha Plan-Apochromat oil immersion objectives (NA 1.46). Environmental conditions were maintained according to each instrument’s specifications, and the same acquisition parameters were applied throughout for data consistency.

For mature mycelium imaging, 5-6-day-old cultures of *M. oryzae* relevant strains were grown on CM 3% agar at 25 °C under a 12 h light/dark cycle. Agar blocks were inverted and mounted on VWR borosilicate cover glasses (22 x 50 mm, thickness no. 1.5) for imaging with a Leica TCS SP8X inverted confocal microscope, using a 63x Plan-Apochromat oil immersion objective (NA 1.4). Z-stacks and time lapse images were captured with combinations of fluorescence channels (488 nm for eGFP via argon laser; 561 nm for TagRFP via DPSS laser) and transmitted light for brightfield imaging, utilizing hybrid detectors (HyD). All acquisition parameters were kept consistent among experimental groups.

For structured illumination microscopy (SIM), conidia of *M. oryzae* relevant strains were prepared on Zeiss high-performance cover glasses (18 x 18 mm, thickness no. 1.5) and mounted similarly. Imaging at 4 and 24 h was performed using a Zeiss Elyra 7 inverted microscope with dual sCMOS cameras for simultaneous two-colour acquisition. Z-stacks were collected with OPSL lasers for 488 nm (eGFP) and 561 nm (TagRFP). A 63x alpha Plan-Apochromat oil immersion objective (NA 1.46) was used. TetraSpeck™ (Thermo Fischer Scientific) 100 nm microspheres served as fiducials to prevent channel misalignment, and image registration was performed using the ZEN Black software (Carl Zeiss AG) channel alignment feature to ensure accurate spatial overlay. Each z-stack comprised 15 phase images per focal plane in Lattice SIM mode. Live-cell imaging was conducted in a chamber with regulated atmosphere (30°C, 5% CO₂, controlled humidity).

#### Large scale *in vitro* appressorium induction for proteomics and Y2H

Conidia were harvested from 10-day-old Petri dish cultures grown on CM agar as described above. Conidial suspensions were inoculated into square Petri plates (12 cm × 12 cm × 1.7 cm) (Greiner Bio-One) to which ten borosilicate cover glasses (Menzel-Gläser, Fisher Scientific UK Ltd.) were adhered using Super Glue (Loctite). Appressorium formation was observed using a Will-Wetzlar inverted light microscope (Wilovert, Hund Wetzlar) to ensure uniform and synchronized development of infection structures. Samples were collected by scraping the cover glass surfaces with a sterile razor blade (Fisher Scientific). The harvested material was snap-frozen in liquid N_2_ and stored at −80°C for subsequent protein extraction.

#### Protein extraction and immunoprecipitation

Lyophilized samples were resuspended in extraction buffer [10 mM dithiothreitol, 1× protease inhibitor cocktail (Sigma-Aldrich), 0.1% IGEPAL, 1 g polyvinylpyrrolidone, 50 ml GTEN (10% glycerol, 25 mM Tris pH 7.5, 1 mM EDTA, 150 mM NaCl)] and mechanically disrupted using a GenoGrinder 2010 (Thermo Scientific) under cold conditions (2 × 1 min at 1,300 rpm). Homogenates were centrifuged (13,200 rpm, 10 min, 4°C; Eppendorf 5415D microcentrifuge), and supernatants transferred to a 5 ml Eppendorf tube. Pellets were re-extracted in fresh extraction buffer, re-disrupted, and centrifuged as above; supernatants were then pooled. Total protein concentration was determined using a Bradford assay (Bio-Rad). Input samples (20 µg protein) were analysed by SDS-PAGE/Western blot after mixing with loading buffer (4× SDS, 1 M DTT). For immunoprecipitation, the remaining total protein extract (∼1.9 ml per sample) was incubated with GFP-Trap beads (Chromotek; 50 µl beads in 100 µl IP buffer [10% glycerol, 25 mM Tris pH 7.5, 1 mM EDTA, 150 mM NaCl, 0.1% Tween 20 (Sigma-Aldrich)]) on a rotary mixer at 4°C for 3 h. Beads were pelleted (800 × g, 1 min), supernatants discarded, and beads washed four times in 1 ml IP buffer with centrifugation as above. Bound proteins were eluted in 40 µl loading buffer (4× SDS, 1 M DTT, H₂O) for 10 min at 70°C, beads pelleted, and 4 µl supernatant analysed by SDS-PAGE/Western blot. ToxA::GFP^51^ and Sep3-6::GFP proteins were detected using a mouse monoclonal GFP antibody (B-2) (Santa Cruz Biotechnology) overnight, followed by washes in 4× TBST (100 ml TBS, 900 ml Milli-Q water, 1 ml Tween 20). Blots were developed using the Thermo Scientific Pierce Fast Western Blot Kit with SuperSignal West Femto Substrate and imaged using an ImageQuant LAS 4000 (GE Healthcare Life Sciences).

#### Sample preparation for LC-MS/MS analysis

Affinity-purified protein samples were fractionated for 1 cm into a 4-20% Mini-PROTEAN TGX Precast gel (Bio-Rad), stained with SimplyBlue SafeStain (Invitrogen), and the resolving portion excised. Gel bands were cut into ∼1 mm³ pieces and destained three times in 50 mM ammonium bicarbonate (ABC)/50% (v/v) acetonitrile (30 min each), dehydrated in 100% acetonitrile (10 min), reduced with 10 mM DTT (30 min, 45°C), alkylated with 55 mM chloroacetamide (20 min, room temperature), and washed three times in 50% (v/v) acetonitrile/ABC (30 min each). Gel pieces were dehydrated again in acetonitrile and rehydrated in trypsin working solution (Pierce Trypsin Protease, MS Grade, cat. no. 90058; 100 ng trypsin in 50 mM ABC, 5% [v/v] acetonitrile; 40 µl per sample). Where required, gel pieces were overlaid with 50 mM ABC, then digested overnight at 37°C. Tryptic peptides were extracted three times with an equal volume of 50% (v/v) acetonitrile, 5% (v/v) formic acid (Pierce LC-MS Grade, cat. no. 85178; 30 min each). Extracts were dried in a SpeedVac, resuspended in 2% (v/v) acetonitrile, 0.2% (v/v) trifluoroacetic acid (Merck, cat. no. 302031) and stored at -20°C. Three to four biological replicates per sample type were analysed.

#### LC-MS/MS analysis

Approximately 35% of each sample was analysed using an Orbitrap Fusion Tribrid Mass Spectrometer coupled online to a Dionex Ultimate 3000 nano-UPLC system (both Thermo Fisher Scientific). Dissolved peptides were injected onto a NanoEase MZ Symmetry C18 trap column (5 µm, 180 µm × 20 mm; Waters) at 20 µl/min in 2% (v/v) acetonitrile, 0.05% (v/v) TFA. Peptides were eluted onto the analytical NanoEase MZ HSS T3 C18 column (1.8 µm, 75 µm × 250 mm; Waters), equilibrated at 3% buffer B (80% acetonitrile, 0.05% formic acid; buffer A, 0.1% formic acid). Elution followed a linear gradient (200 nl/min): 2.5 min 3% B, 5 min 6.3% B, 13 min 12.5% B, 50 min 42.5% B, 58 min 50% B, 61 min 65% B, 63-66 min 99% B, 67-90 min 3% B. The mass spectrometer operated in positive ion mode (nano-electrospray, +2.2 kV; PicoTip emitter 10 µm ID [New Objective]; capillary temp. 275°C). Data-dependent acquisition used full MS scans (m/z 300-1,800; resolution 120,000; AGC target 1 × 10⁶) followed by MS/MS on the 40 most abundant precursors (charge states 2⁺-7⁺; resolution 120,000; AGC target 1 × 10⁵; isolation width 1.6 m/z; NCE 30%; dynamic exclusion 30 s; minimum AGC 5 × 10³; intensity threshold 4.8 × 10⁴). Peptide matching and isotope exclusion were enabled.

#### RNA extraction and cDNA synthesis

Total RNA was extracted from ground mycelial tissue after 48 h growth in liquid CM medium and from spores undergoing appressorium development at 0, 2, 4, 8, and 16 h. RNA was purified using the RNeasy Plant Mini Kit (Qiagen) according to the manufacturer’s instructions. First-strand cDNA was synthesized from extracted RNA using the GoScript™ Reverse Transcription System (Promega) following the manufacturer’s protocol.

#### Hybrigenics Yeast Two-Hybrid based protein interactions screening

A 1 mg aliquot of total RNA pooled from *M. oryzae* spores undergoing appressorium development at 0, 2, 4, 8, and 16 h was provided to Hybrigenics (Paris, France) for construction of the ULTImate Y2H™ cDNA library (MAOR_RP). From this RNA, mRNA was isolated (GenElute™ mRNA Miniprep Kit, Sigma-Aldrich; yielding 6.5 µg polyA⁺ RNA), random-primed double-stranded cDNA synthesized (SuperScript®, Invitrogen; average insert 500-700 nt), cloned into the pP6 prey vector, and transformed into *Escherichia coli* (50 million independent clones) and yeast (10 million clones). Library quality was confirmed by sequencing 192 random clones (7% empty/contaminated vectors).

Core septins served as bait constructs in Hybrigenics’ high-throughput mating platform:

- Sep3 (G4MTD7, aa 1-443; pB66; 49.3 million interactions tested; 0 mM 3-AT)
- Sep4 (G4ML89, aa 1-337; pB66/pB35;122 million interactions; 0-0.5 mM 3-AT)
- Sep5 (G4NKC0, aa 1-383; pB66; 80.5 million interactions; 0.5 mM 3-AT)
- Sep6 (G4N184, aa 1-385; pB35; 50.5 million interactions; 0 mM 3-AT)

Positive diploids were selected on histidine-deficient media and prey fragments sequenced/mapped to the *M. oryzae* genome (Ensembl Fungi, MGv8). Bioinformatics analysis (Predicted Biological Score, PBS) was used to rank confidence in the interactions.

#### Yeast-Two-Hybrid (Y2H) analysis

Protein-protein interactions between core septins (Sep3-Sep6) and five candidates identified by IP-MS and Hybrigenics yeast two-hybrid screening were assessed using the Matchmaker™ Gold Yeast Two-Hybrid System (Takara Bio). Bait (pGBKT7) and prey (pGADT7) constructs were co-transformed into chemically competent *Saccharomyces cerevisiae* Y2HGold cells (Takara Bio) using the Frozen-EZ Yeast Transformation II™ kit (Zymo Research) per manufacturer’s instructions. Transformants were selected on SD/-Leu/-Trp medium. Single colonies were inoculated into 5 ml SD/-Leu/-Trp, grown overnight at 30°C, normalized to OD₆₀₀ = 1.0, and serially diluted (10⁰-10⁻³). Aliquots (5 µl) from each dilution were spotted onto SD/-Leu/-Trp (growth control), SD/-Leu/-Trp/-His (low stringency), and SD/-Leu/-Trp/-His/-Ade + X-α-Gal + aureobasidin A (high stringency). Plates were incubated at 30°C for 60-72 h prior to imaging.

#### Generation of Δ*msi1* deletion mutant

DNA sequence for *MSI1* (MGG_01508) was retrieved from the *M. oryzae* genome database (http://fungi.ensembl.org/Magnaporthe_oryzae/Info/Index) and used to design primers. PCR fragments were amplified from genomic DNA and plasmid templates using primers listed in Table S9. The Δ*msi1* deletion cassette was generated using a split-marker strategy with hygromycin B resistance as the selectable marker.^52^ The two linear fragments were co-transformed into Guy11 protoplasts to promote homologous recombination. Transformants were selected on Hygromycin B and verified by whole-genome sequencing (Figure S4C). A representative Δ*msi1* mutant was retained for subsequent analyses.

#### Generation of fluorescently tagged strains

DNA sequences of genes of interest were retrieved from the *Magnaporthe oryzae* genome database (Ensembl Fungi; http://fungi.ensembl.org/Magnaporthe_oryzae) and used for primer design (Table S9). PCR fragments were amplified from genomic DNA or plasmid templates and cloned together with GFP or TagRFP into pCB1532-SUR (for GFP) or pScBAR (for TagRFP) using the In-Fusion™ HD Cloning Kit (Takara Bio). Primers contained overhangs homologous to vector sequences to facilitate seamless assembly. Constructs were sequence-verified by Sanger sequencing and transformed into respective *M. oryzae* protoplasts for ectopic integration. Transformants were selected on sulfonylurea or glufosinate ammonium and verified by diagnostic PCR; a representative transformant of each strain was used for live-cell imaging.

Septin complementation strains expressing individual GFP-tagged core septins (Sep3, Sep4, Sep5, and Sep6) were previously reported.^24^ The Sep4-GFP/Sep5-TagRFP double-labeled strain was generated by sequential transformation. Briefly, a SEP4-GFP translational gene fusion was cloned into linearized pCB1532-SUR, sequence-verified, and introduced into Δ*sep4*.^24^ The SEP5-TagRFP construct was subsequently cloned into pScBAR and transformed into the Δ*sep4* Sep4-GFP background strain. Msi1-5-GFP strains were created by cloning *MSI1*-*5* open reading frames translationally fused to GFP into linearized pSB1532-SUR using In-Fusion™ HD Cloning (Takara Bio). After sequence verification, constructs were transformed into *M. oryzae* Δ*sep4* Sep4-TagRFP and Δ*sep6* Sep6-TagRFP background strains (Table S10). The Δ*msi1* complementation strain was obtained by reintroducing the *MSI1*-GFP construct into the Δ*msi1* mutant background. For competitive pathogenic fitness assays,^38^ a TrpC::mChFP plasmid was transformed into Guy11 (yielding WT-mCherry), and a TrpC::GFP plasmid was introduced into the Δ*msi1* mutant (Δ*msi1*-GFP) as described in Table S10.

#### Y2H plasmid construction

Coding sequences of the core septins (Sep3-Sep6) were cloned into pGADT7 (prey) and Msi1-Msi5 into pGBKT7 (bait) using In-Fusion™ HD Cloning (Takara Bio). PCR fragments were amplified from *M. oryzae* cDNA prepared from mycelia grown in liquid complete medium (CM) using primers containing 15-bp homologous overhangs matching BamHI/EcoRI-linearized vectors (Table S10). All constructs were verified by Sanger sequencing prior to transformation into yeast.

#### Transformation of *M. oryzae*

*M. oryzae* protoplasts were prepared and transformed using a polyethylene glycol (PEG)-mediated transformation protocol. The actively growing colony margin was excised from a CM plate and blended in 150 ml liquid CM. The culture was incubated at 25°C with shaking (100 rpm) for 48 h, harvested by filtration through sterile Miracloth (Millipore), and the mycelium washed with sterile water. Mycelium was transferred to a 50 ml Falcon tube and resuspended in 25 ml sterile-filtered 0.7 M NaCl containing 250 mg lysing enzymes from *Trichoderma harzianum*. The suspension was gently dispersed to break hyphal clumps, wrapped in aluminum foil, and incubated at 30°C with shaking (75 rpm) for 3-4 h. Protoplasts were filtered through sterile Miracloth (Millipore), pelleted by centrifugation (3,000 rpm, 15 min, 4°C; Beckman JS-13.1 rotor, J2.MC centrifuge), washed twice in STC buffer (1.2 M sucrose, 10 mM Tris-HCl pH 7.5, 10 mM CaCl₂), and finally resuspended in 500 µl STC (counted using a Neubauer haemocytometer). Transformation was performed with 5 × 10⁶ protoplasts mixed with 6-10 µg DNA in STC and incubated at room temperature for 30 min. The mixture was then treated with 1 ml PTC solution (60% PEG4000, 10 mM Tris-HCl pH 7.5, 10 mM CaCl₂) added in aliquots with gentle inversion (7 times), followed by a 15 min incubation at room temperature. The transformation mixture was transferred using a wide-bore pipette tip to 5 ml liquid OCM (for HygB selection: 50 ml/L 20× nitrate salts, 1 ml/L 1000× trace elements, 1 ml/L vitamin solution, 10 g/L D-glucose, 2 g/L tryptone, 1 g/L yeast extract, 1 g/L casein hydrolysate, 273.8 g/L 0.8 M sucrose; pH 6.5) or BDCM (for BAR/SUR selection: 1.7g/L yeast nitrogen base without amino acids/ammonium sulfate [BD Difco], 2 g/L NH₄NO₃, 1 g/L L-asparagine, 10 g/L glucose, 273 g/L sucrose; pH 6.0) in a foil-wrapped 50 ml Falcon tube. Cultures were incubated overnight at 25°C with shaking (75 rpm; ∼16 h). The mixture was added to 150 ml molten bottom agar (BDCM for BAR/SUR or OCM for HygB, each supplemented with 15 g/L agar) and poured into 5 sterile Petri dishes (∼20-25 ml per plate). Plates were incubated in the dark at room temperature for ≥16 h, then overlaid with ∼15 ml molten top agar containing selective agents: CM (250 µg/ml hygromycin B), BDCM (150 µg/ml BAR or 300 µg/ml SUR)

#### Fungal genomic DNA extraction

*M. oryzae* genomic DNA was extracted using the DNeasy Plant Mini Kit (Qiagen). Mycelium was grown in liquid CM by harvesting spores from a CM plate and inoculating a 150 ml flask of liquid CM covered with a foam stopper and sterilized aluminum foil. The flask was incubated on a rotary shaker at 100 rpm and 25°C with a 12 h photoperiod for 48 h. Cultures were harvested by filtration through sterile Miracloth and mycelia were washed with sterile water. Mycelial samples were disrupted using a mortar and pestle in liquid N_2_. To each ground sample, 400 µl Buffer AP1 and 4 µl RNase A were added, followed by thoroughly mixing and incubation for 10 min at 65°C, with inversion of the tube 2-3 times during incubation. Subsequently, 130 µl Buffer P3 was added, mixed, and the samples were incubated for 5 min on ice. The lysate was centrifuged for 5 min at 20,000 × g (14,000 rpm), and the supernatant was transferred to a QIAshredder spin column placed in a 2 ml collection tube and centrifuged for 2 min at 20,000 × g. The flow-through was transferred to a new tube without disturbing any pellet. One and a half volumes of Buffer AW1 were added and mixed by pipetting. Aliquots of 650 µl were loaded onto a DNeasy Mini spin column in a 2 ml collection tube and centrifuged for 1 min at ≥6,000 × g (≥8,000 rpm); the flow-through was discarded and the step was repeated with the remaining sample. The spin column was then placed into a new 2 ml collection tube, 500 µl Buffer AW2 was added, and the column was centrifuged for 1 min at ≥6,000 × g. After discarding the flow-through, an additional 500 µl Buffer AW2 was added and centrifuged for 2 min at 20,000 × g. The spin column was transferred to a new 1.5 or 2 ml microcentrifuge tube, 50 µl double distilled H₂O was added for elution, and the column was incubated for 5 min at room temperature (15-25°C) before centrifugation for 1 min at ≥6,000 × g. The elution step was repeated.

For rapid DNA extraction, a 1-5 mm² piece of mycelium was removed from the growing margin of a 5-day-old *M. oryzae* culture using a sterile steel scalpel and transferred to a 2 ml microcentrifuge tube containing one sterile metal homogenizer bead. Then, 500 µl Breaking buffer (4 ml EDTA 0.5 M, 20 ml Tris 1M pH 7.5, 14.9 g KCl, ad 200 ml double distilled H_2_O) was added to the sample. Mycelium was disrupted in a Geno/Grinder for 2 min at 1,300 rpm at room temperature (RT), followed by centrifugation for 10 min at 16,000 × g at RT. A 300 µl aliquot of the supernatant was transferred to a fresh 1.5 ml microcentrifuge tube, 300 µl isopropanol was added, and the tube was inverted several times before incubation at 4°C overnight. Samples were centrifuged for 10 min at 16,000 × g at RT, the supernatant was discarded, and the tube was placed upside down with the lid open on tissue paper for 10 min to air-dry the pellet. The samples were then dried in a SpeedVac for 10 min at 30°C. Finally, 50 µl double distilled H₂O was added, and the tube was incubated for 15 min at 37°C.

#### Gene deletion confirmation by whole genome sequence analysis (WGS)

*M. oryzae* genomic DNA was purified and quantified using a NanoDrop spectrophotometer (Thermo Scientific). Paired-end sequencing was performed by Novogene (Cambridge, UK) on an Illumina NovaSeq 6000 system (2 lanes per sample). The *M. oryzae* reference genome (MGv8 assembly) was indexed with Bowtie2 v2.2.5,^53^ and sequencing reads from Δ*msi1* mutants were aligned to generate SAM files. Alignments were converted to coordinate-sorted BAM files using SAMtools v0.1.9,^54,55^ indexed, and visualized in IGV v2.16.2 ^56,57^ alongside MGv8 reference genome and annotations. Targeted gene deletion was confirmed by complete absence of read coverage across the *MSI1* locus (MGG_01508) using IGV v2.16.2. (Figure S4C).^56,57^

#### Rice Blast Disease Assays

Detached leaf infection assays were performed on barley (*Hordeum vulgare* cv. Golden Promise). The first leaf of 1-week-old seedlings was excised, placed on filter paper in square Petri dishes, and secured with autoclave tape. Filter paper was moistened with water and both cut leaf ends were covered with wet cotton pads before sealing the dishes. Conidial suspensions were prepared from 10-day-old CM plates and adjusted to 1 × 10⁴ spores/ ml in double distilled H₂O. Drops of 5 µl were applied to the leaf surface. Inoculated leaves were incubated in a controlled growth room at 25°C, with the dish lids misted with water to maintain humidity. At 5 days post inoculation (dpi), leaves were collected, mounted on Whatman paper, and imaged; disease lesion size was quantified using Fiji (ImageJ2 Version:2.14.0/1.54f).

Attached leaf spray infection assays were performed on intact 3-week-old rice (*Oryza sativa* cv. CO-39) seedlings. Conidial suspensions were prepared from 10-day-old CM plates and adjusted to 1 × 10⁵ spores/ml in 0.2% (w/v) gelatin in double distilled H₂O. Two pots of plants per fungal isolate were placed in a black tray containing wet tissue paper, enclosed in a large polyethylene bag, and sprayed with 5 ml conidial suspension using a small mist sprayer (Ampulla LTD) to ensure a fine, uniform coating of the leaf surface. Inoculated plants were maintained in the sealed bag and incubated in a growth cabinet at 26°C (10 h light) / 24°C (14 h dark) for 5 days. Infected leaf tips (∼5 cm) were then excised, attached to Whatman paper with adhesive tape, and blast lesions were imaged and assessed at 5 dpi.

For leaf sheath infection assays, conidial suspensions were prepared from 10-day-old CM plates and adjusted to 1 × 10³ spores/ml in 0.2% (w/v) gelatin. Leaf sheaths were collected from 4-week-old rice plants, the outermost and innermost layers removed, and the remaining sheath cut into 7-8 cm sections. Sheaths were placed horizontally on two eight-tube PCR strips in square Petri dishes lined with moist paper towels and inoculated by filling the interior with spore suspension using a 200 µl pipette. Inoculated sheaths were incubated at 25°C. For confocal microscopy, ultrathin sections (3-4 cell layers) were cut with a sharp blade, excess outer epidermis of the midvein carefully shaved away, remaining transparent cell layers trimmed, and mounted in water on glass slides under 22 × 50 mm coverslips. Cell-to-cell invasion by invasive hyphae was quantified at 48 hpi across infection stages (appressoria without penetration hyphae, first cell invasion, second cell invasion), scoring 100 infection sites per experiment per mutant strain relative to Guy11 wild type.

#### Competition pathogenicity fitness assay

Relative fitness assays were performed using a Guy11 strain expressing cytoplasmic mChFP under the TrpC constitutive promoter (WT-mCherry) and the Δ*msi1* mutant expressing cytoplasmic GFP under the same TrpC promoter (Δ*msi1*-GFP) [Yan et al., 2023]. Conidia from both strains were mixed in a 1:1 ratio and used to inoculate Golden Promise barley seedlings via detached leaf infection assays. Blast disease was allowed to progress until necrotic lesions formed, from which conidia were recovered and their WT-mCherry:Δ*msi1*-GFP ratios quantified by fluorescence. Recovered conidia were prepared at their new proportions to inoculate fresh barley seedlings, and the competitive cycle was repeated for three generations. The Δ*msi1* TrpC::GFP mutant was driven to near extinction by generation three. Data are presented as the proportion of each strain recovered per generation (mean ± SE, n = 3 biological replicates) and relative fitness calculated as x₂(1 − x₁)/x₁(1 − x₂), where x₁ is the initial frequency and x₂ is the final frequency of the test strain conidia (Figure 6F).

#### Spore germination frequency assays

Conidia were harvested from 10-day-old CM plates, filtered through Miracloth, and adjusted to 1 × 10⁵ spores/ml in double distilled H₂O. Aliquots (30 µl) were spread onto glass coverslips placed in square Petri dishes above wet tissue paper to maintain humidity and incubated at 25°C. Germination was scored microscopically at 2 hpi by counting ≥100 conidia per replicate. The frequency of germination (%) was calculated as (germinated spores/total spores) × 100. Three biological replicates with three technical replicates each were performed.

#### Fungal colony growth assays

Colony growth assays were performed on both minimal medium (MM) and complete medium (CM). For MM, 5 mm round agar plugs from the edge of actively growing colonies of Guy11 WT, Δ*msi1*, and complementation strains were punched out and transferred to MM plates, which were incubated at 25°C for 14 days. For CM, plugs were transferred to CM plates and incubated at 18°C, 25°C, and 30°C for 7 days. Area of growth (mm²) was quantified using Fiji from images captured at the end of each incubation period. Box plots represent one representative replicate from three independent biological replicates (n = 3), with significant differences between strains determined by Student’s t-test (p ≤ 0.05, marked by asterisks).

#### Creation of Schematics

The schematic illustrations in this article were created with BioRender.

### QUANTIFICATION AND STATISTICAL ANALYSIS

#### Data processing

Raw files (.raw format) were processed using MaxQuant (*v* 2.4.1.0),^58^ with default parameters unless specified otherwise. MS/MS spectra were searched against *Magnaporthe oryzae* reference proteome (MG8, downloaded from Ensembl Fungi https://fungi.ensembl.org/Magnaporthe_oryzae/Info/Index)^59^ along with in-house constructs and contaminants database. Protein database searches were performed using the integrated Andromeda peptide search engine.^60^ Search parameters involved trypsin/P as proteolytic enzyme, allowing cleavage after lysine and arginine residues, including cases where proline follows the cleavage site, with a maximum of two missed cleavages allowed to account for incomplete digestion. Fixed modifications were set to carbamidomethylation of cysteine residues, to account for the alkylation step during sample preparation. Oxidation of methionine and acetylation of protein N-termini were set as variable modifications, to capture common post-translational modifications. Precursor ion mass tolerance was set to 20 ppm for the first search, allowing initial mass calibration and then narrowed to 6 ppm for the main search. Fragment ion tolerance was set to 20 ppm for high-resolution MS/MS spectra on an Orbitrap analyzer. False discovery rate (FDR) was controlled at 5% (q-value < 0.05) at both the peptide spectrum match (PSM) and protein group levels using a target-decoy approach with reversed protein sequences.^61^ Label-Free Quantification (LFQ) was used to quantify relative protein abundances across samples ^62^ with FastLFQ algorithm, which uses an accurate and computationally fast normalization procedure for LFQ intensity calculation.^63^ The match between runs feature was enabled with a matching time window of 1.0 minute, allowing the transfer of peptide identifications between runs based on their mass and retention times.^62^ Protein quantification was based on unique and razor peptides. For a protein to be identified, at least one peptide was required. For a protein to receive a quantitative intensity value, at least two peptide measurements across samples were required, ensuring reliable abundance estimates.

#### Detection of putative septin interactors

MaxQuant output files (proteinGroups.txt) were imported into R (v4.3.1) for downstream processing and quality control. Potential contaminants and reverse database hits were filtered out from the dataset. Putative septin interactors were identified by calculating fold changes for each protein as the ratio of the mean LFQ intensity across biological replicates in septin immunoprecipitates to mean LFQ intensity across biological replicates in pToxA-GFP controls at each corresponding timepoint. A protein was qualified as a putative septin interactor if it was quantified in at least two biological replicates for a given septin-timepoint combination, and it had at least 2-fold intensity relative to pToxA-GFP control in at least one timepoint. Statistically significant interactors based on a bootstrap t-test (p ≤ 0.05) were defined as high-confidence septin-interactors. The complete dataset of putative septin interactors, including LFQ intensities, fold-changes, and associated metadata, is provided in Table S1, while high-confidence interactors are provided in Table S2. Putative septin interactors were stratified into fold-change categories (FC>=2, FC>10, FC>100 and FC>1000) to visualize the strength of septin associations (Figure 2 A, Figure S2B). To identify shared and unique septin associations, binary presence–absence matrices were generated for each protein across the four septins. The number of septin combinations in which each protein was detected was used to calculate set intersections using the UpSetR package,^64^ followed by visualisation of overlapping and unique interactors across the four core septins during developmental time points (Figure 2B), mycelium (Figure S2C), and yeast two-hybrid data (Figure 5C).

#### Network analyses of septin interactome

Putative septin interactors from the four septins across the six developmental timepoints (0h, 2h, 4h, 8h, 16h and 24h) and their UpSet combinations were used to construct a global septin interactome network. Network data were processed and generated using R (v4.3.1) using a custom script and rgexf package.^65^ The resulting GEXF (graph exchange XML format) file was exported for network visualization in Gephi.^66^ The exported network was structured as an undirected graph consisting of 938 nodes representing individual proteins (septin interactors), and 1592 edges representing co-occurrence of proteins within the same interaction set. Network layout was directed using ForceAltas2 layout algorithm,^67^ with parameters optimized to balance network readability (scaling =2.0, gravity 1.0, prevent overlap enabled). Nodes were labelled as gene names retrieved from MagnaGenes v.1.0^37^ where available, whereas interactors with no annotated gene names were labeled with MGG identifiers, however label visibility was selectively reduced to improve readability. Nodes colors were based according to their Upset combinations as provided in Figure 2B. Temporal dynamics of the septin interactome were visualized by generating time-resolved network snapshots at each developmental stage (0-24h) (Figure 2D). More broader annotations based on their experimentally validated roles in biological processes, using literature curated data from MagnaGenes v.1.0^37^ was used to visualize time-resolved networks related to these processes.

#### Gene function enrichment of putative septin interactors

To visualize the temporal dynamics of putative septin interactors and identify timepoint specific functional modules, normalized LFQ intensities for the four septins (Sep3, Sep4, Sep5, Sep6) across six developmental timepoints (0h, 2h, 4h, 8h, 16h, 24h) were used to calculate log2-transformed fold changes relative to ToxA controls. For each interactor, the timepoint showing maximum fold change was used to group the interactors into temporal modules. Fold-changes were normalized into z-scores and visualised using heatmaps constructed using ComplexHeatmap package in R.^68^ Proteins were further arranged into three temporal sectors - early (0h, 2h), mid (4h, 8h) and late (16h, 24h) stages. Heatmaps were clustered based on hierarchical clustering (complete linkage, Minkowski distance). Gene Ontologies (GO), InterPro domains ^69^ and other functional annotations for *M. oryzae* were downloaded from Ensembl Fungi ^59^ through the Biomart portal.^70^ GO enrichment analysis was performed for each septin-timepoint combination using the topGO package ^71^ in R with the weight01 algorithm and Fisher’s exact test, to calculate the significantly enriched (p ≤ 0.05) biological processes, molecular functions and cellular components. Representative significantly enriched GO terms (Table S4 and 5) were selected and shown on the heatmaps, with term selection based on known and novel roles in interaction with septins. A detailed overview of all 1179 putative septin interactors across all four core septins and all the developmental timepoints including the mycelium stage, were visualized as a heatmap using ComplexHeatmap (Figure S1).^68^ Proteins in this global heatmap were ordered primarily by their highest fold change relative to ToxA controls across all for septins, and secondarily clustered using hierarchical clustering (average linkage, Pearson’s distance) to group proteins with similar temporal patterns.

#### Processes-level enrichment of the septin interactome

To identify developmental processes significantly enriched among putative septin interactors, Fisher’s exact test was performed for each septin timepoint combination using annotations based on literature-curated data from MagnaGenes v.1.0.^37^ For each process, a 2×2 contingency table was calculated, where A = interactors in the process, B = interactors not in the process, C = non-interactors in the process and D = non-interactors in the process with the entire *M. oryzae* proteome as background. Fisher’s exact test was performed using fisher.test function in R, resulting in two-tailed p-values and odds ratios to assess the over- or under-representation. Multiple testing adjustment for -p-values was done using the Benjamini-Hochberg FDR correction,^72^ with a significance threshold of FDR ≤ 0.05. Results were visualized as interactor-to-category ratios in heatmaps using pheatmap package in R.^73^

#### Conversation analysis of BAR domain-containing proteins

To identify evolutionary conservation and copy number variation of BAR domain-containing proteins in other fungi, we performed a comparative analysis using proteomes of 41 fungal species. The list of species with their respective associated lifestyles (biotrophic, necrotrophic, hemibiotrophic, saprophytic) were obtained from a published dataset.^74^ Proteomes for all 41 species were downloaded from publicly available databases, and FASTA headers were standardized to maintain consistency throughout the analyses. Orthologous proteins were grouped using OrthoFinder v2.5.4 with default parameters.^75^ BAR domain proteins were identified by querying the previously mentioned InterPro annotations using the identifiers IPR027267, IPR031160, IPR039463, IPR004148, IPR037429, IPR035803, IPR018859, IPR045734. Copy numbers of BAR domain-containing proteins were retrieved from the Orthogroups file and plotted along the species tree generated by OrthoFinder (Figure S4A).

#### Statistical analysis

Leaf-drop virulence assays on barley (cv. Golden Promise), colony growth on minimal medium (MM) agar plates, and conidial germination rates were analysed using ordinary one-way ANOVA. The effect of temperature (18, 25 and 30°C) on colony growth on complete medium (CM) agar plates was analysed using two-way ANOVA with strain and temperature as factors. In planta penetration and invasive growth on rice (cv. CO-39) were analysed using unpaired t-tests. The competitive fitness assay was analysed using multiple paired t-tests. Data were obtained from ≥3 independent biological replicates and are presented as mean ± SE. Differences were considered statistically significant at p ≤ 0.05. GraphPad Prism 10.4.1 (GraphPad Software Inc.; Dotmatics) was used for all analyses.

#### Use of AI tools

AI-assisted tools (ChatGPT, OpenAI) were used to improve the clarity and language of the manuscript. These tools were not used for data analysis, figure generation, or scientific interpretation. All results and conclusions were generated and verified by the authors.

---

**Figure S1. Hierarchical clustering heat map of putative septin interactors during appressorium development in *M. oryzae*.**

Heat map to illustrate hierarchical clustering of 1179 putative septin-interacting proteins identified during appressorium development at 0, 2, 4, 8, 16, and 24 h post-inoculation for each septin (Sep3, Sep4, Sep5, and Sep6), based on comparative proteomics workflow detailed in Figure 1C. Heat map is organised into four blocks, corresponding to each septin, with rows representing individual septin-protein interaction measured across the six time points. Protein fold changes (FC) relative to a control (toxA-GFP) shown using a colour gradient from blue (low FC) to red (high FC), representing z-scores-normalised fold changes derived from immunoprecipitation mass spectrometry (IP-MS). Asterisks indicate time point at which a protein met stringent septin-interaction criteria (fold change ≥ 2 and detection in at least two biological replicates). Proteins are ordered using hierarchical clustering based on similarity in FC profiles further arranged by peak FC time point (indicated on the left), revealing distinct temporal interaction patterns within the septin network. Protein identifiers (MGG IDs) and functional annotations are shown on the right.

**Figure S2. Dynamic localisation and comparative proteomic analysis reveal a conserved set of septin interactors during vegetative growth of *M. oryzae.***

(A) Confocal laser scanning microscopy reveals dynamic localisation of the four septins (Sep3-GFP, Sep4-GFP, Sep5-GFP, and Sep6-GFP) during vegetative growth. Representative images show septin localisation at hyphal tip, septum, and branch (left to right). Scale bars: 5 μm.

(B) Quantitative summary of putative septin interactors identified for Sep3, Sep4, Sep5, and Sep6 during vegetative growth. Bar heights indicate total number of interactors per septin, with segmented shading representing different fold-change thresholds (FC ≥ 2, > 10, > 100, and > 1000). The total number of interactors is shown above each bar.

(C) UpSet plot illustrating shared and unique septin interactors during vegetative growth, organised into four major categories: (I) interactors associated with a single septin, (II) interactors shared by two septins, (III) interactors shared by three septins, and (IV) interactors common to all four septins. Lines connecting septin nodes below the bars indicate specific septin combinations for each group of interactors. The fifteen interaction patterns shown include Sep3; Sep4; Sep5; Sep6; Sep3-Sep4; Sep3-Sep5; Sep3-Sep6; Sep4-Sep5; Sep4-Sep6; Sep5-Sep6; Sep3-Sep4-Sep5; Sep3-Sep4-Sep6; Sep4-Sep5-Sep6; Sep3-Sep5-Sep6; and Sep3-Sep4-Sep5-Sep6, each distinguished by a unique colour.

(D) Venn diagrams showing shared and unique septin interactors of Sep3, Sep4, Sep5 and Sep6 identified during appressorium development and vegetative growth. Each diagram illustrates interactors unique to appressorium development, unique to vegetative growth, and those common to both conditions.

**Figure S3. Dynamic spatial association of candidate septin interactors with septins during appressorium development and vegetative growth of *M. oryzae.***

(A) Confocal laser scanning microscopy combined with Airyscan super-resolution imaging reveals dynamic localisation of four *<u>M</u>agnaporthe* <u>s</u>eptin <u>i</u>nteractors (Msi1-GFP, Msi2-GFP, Msi3-GFP, and Msi5-GFP) during appressorium development at 0, 2, 4, 8, 16, and 24 hours post inoculation. Scale bars: 10 μm. Co-expression of Msi-GFP labelled proteins with Sep4-TagRFP is shown. Asterisks indicate time points at which immunoprecipitation mass spectrometry (IP-MS) detected interactions with core septins.

(B) Confocal laser scanning microscopy analysis of candidate interactor Msi4 tagged with GFP and co-expressed with Sep6-TagRFP during vegetative growth. Images (top to bottom) show mid-plane sections of Msi4-GFP, Sep6-TagRFP, and their merged overlay. Scale bar: 5 μm.

**Figure S4. Functional characterisation of the BAR domain protein Msi1 in *M. oryzae* reveals evolutionary conservation of fungal BAR domain proteins and suggests a role *in planta*.**

(A) Phylogenetic analysis of BAR domain-containing proteins across 42 fungal species representing diverse lifestyles (biotrophic, necrotrophic, hemibiotrophic, saprophytic) to assess evolutionary conservation and copy number variation. Purple boxes surrounding MGG identifiers highlight seven septin interacting BAR domain proteins identified in this dataset.

(B) Schematic representation of the *MSI1* gene deletion strategy using homologous recombination via a split marker approach.

(C) Whole genome sequencing analysis of a confirmed Δ*msi1* mutant strain showing loss of *MSI*1 gene coverage due to insertion of the HPH resistance cassette.

(D) Representative colony morphology of wild-type Guy11 and Δ*msi1* strains after 7 days of growth on complete medium.

(E) Quantitative assessment of spore germination rates (%) on coverslips after 2 h incubation at 25°C for Guy11 WT and Δ*msi1* strains. Bars represent mean ± SE from one representative experiment, repeated three times with similar results (Ordinary one-way ANOVA, p ≤ 0.05).

(F) Quantitative analysis of colony area (mm²) on minimal medium plates after 14 days of growth at 25°C, comparing Guy11 WT, Δ*msi1* and complementation strains. Box plots show one representative replicate from three independent experiments, with significant differences indicated by asterisks (Ordinary one-way ANOVA, p ≤ 0.05).

(G) Quantitative analysis of colony area (mm²) on complete medium plates after 7 days of growth at 18 °C, 25°C and 30 °C, comparing Guy11 WT, Δ*msi1* and complementation strains. Box plots show one representative replicate from three independent experiments, with significant differences indicated by asterisks (Two-way ANOVA, p ≤ 0.05).

(H) Confocal laser scanning microscopy of Msi1-GFP localisation during infection (∼36 h post-inoculation) in rice leaf sheath assays using 3-week-old cultivar CO-39 seedlings. Images show infection structures with punctate fluorescence, indicating dynamic subcellular localisation of Msi1 in plant infections. Scale bar: 10μm A supplementary video (Video 1) demonstrates the mobility of Msi1-GFP puncta during infection.

**Supplementary Video 1. Msi1-GFP localises as mobile puncta in invasive hyphae of *M. oryzae* during plant infection.**

Fluorescence microscopy showing Msi1-GFP localization ∼36 h post-inoculation in leaf sheath assays using 3-week-old rice CO-39 seedlings. The video captures infection structures exhibiting punctate fluorescent signals that are highly dynamic and mobile, consistent with subcellular localisation of Msi1 *in planta*. Scale bar: 10 μm; time: seconds).

## Supplementary Tables

**Table S1:** Complete septin interactome dataset during all infection-related developmental time points and in mycelium of *M. oryzae*

**Table S2:** Filtered interactome data containing only statistically significant septin interactors

**Table S3:** MS-based quantification confirming expression and recovery of GFP-tagged septins

**Table S4:** Significantly enriched Gene Ontology terms identified in septin time point combinations during appressorium development

**Table S5:** Significantly enriched Gene Ontology terms for septin mycelium combinations

**Table S6:** Fisher’s exact test results for septin interactome-enriched gene categories from functional categories of the MagnaGenes database

**Table S7:** Yeast two-hybrid (Y2H) interaction data from ultra-high throughput Hybrigenics screening

**Table S8:** Genes identified by both IP-MS and Y2H interaction analyses

**Table S9:** Oligonucleotide primers used in this study

**Table S10:** Key Resources Table

## REFERENCES

1. Delic, S., Shuman, B., Lee, S., Bahmanyar, S., Momany, M., and Onishi, M. (2024). The evolutionary origins and ancestral features of septins. Front Cell Dev Biol 12, 1406966. 10.3389/fcell.2024.1406966.

2. Spiliotis, E.T., and Nakos, K. (2021). Cellular functions of actin- and microtubule-associated septins. Curr Biol 31, R651–r666. 10.1016/j.cub.2021.03.064.

3. Mostowy, S., and Cossart, P. (2012). Septins: the fourth component of the cytoskeleton. Nat Rev Mol Cell Biol 13, 183–194. 10.1038/nrm3284.

4. Weirich, C.S., Erzberger, J.P., and Barral, Y. (2008). The septin family of GTPases: architecture and dynamics. Nat Rev Mol Cell Biol 9, 478–489. 10.1038/nrm2407.

5. Spiliotis, E.T., and McMurray, M.A. (2020). Masters of asymmetry - lessons and perspectives from 50 years of septins. Mol Biol Cell 31, 2289–2297. 10.1091/mbc.E19-11-0648.

6. Van Ngo, H., and Mostowy, S. (2019). Role of septins in microbial infection. J Cell Sci 132. 10.1242/jcs.226266.

7. Dolat, L., Hu, Q., and Spiliotis, E.T. (2014). Septin functions in organ system physiology and pathology. Biol Chem 395, 123–141. 10.1515/hsz-2013-0233.

8. Oh, Y., and Bi, E. (2011). Septin structure and function in yeast and beyond. Trends Cell Biol 21, 141–148. 10.1016/j.tcb.2010.11.006.

9. Weems, A., and McMurray, M. (2017). The step-wise pathway of septin hetero-octamer assembly in budding yeast. Elife 6. 10.7554/eLife.23689.

10. Bridges, A.A., and Gladfelter, A.S. (2015). Septin Form and Function at the Cell Cortex. J Biol Chem 290, 17173–17180. 10.1074/jbc.R114.634444.

11. Woods, B.L., and Gladfelter, A.S. (2021). The state of the septin cytoskeleton from assembly to function. Curr Opin Cell Biol 68, 105–112. 10.1016/j.ceb.2020.10.007.

12. Papachristou, E.K., Kishore, K., Holding, A.N., Harvey, K., Roumeliotis, T.I., Chilamakuri, C.S.R., Omarjee, S., Chia, K.M., Swarbrick, A., Lim, E., et al. (2018). A quantitative mass spectrometry-based approach to monitor the dynamics of endogenous chromatin-associated protein complexes. Nat Commun 9, 2311. 10.1038/s41467-018-04619-5.

13. Hashimoto, Y., Sheng, X., Murray-Nerger, L.A., and Cristea, I.M. (2020). Temporal dynamics of protein complex formation and dissociation during human cytomegalovirus infection. Nat Commun 11, 806. 10.1038/s41467-020-14586-5.

14. Wright, S.N., Colton, S., Schaffer, L.V., Pillich, R.T., Churas, C., Pratt, D., and Ideker, T. (2025). State of the interactomes: an evaluation of molecular networks for generating biological insights. Mol Syst Biol 21, 1–29. 10.1038/s44320-024-00077-y.

15. Wu, S., Zhang, S., Liu, C.M., Fernie, A.R., and Yan, S. (2025). Recent Advances in Mass Spectrometry-Based Protein Interactome Studies. Mol Cell Proteomics 24, 100887. 10.1016/j.mcpro.2024.100887.

16. Fields, S., and Song, O. (1989). A novel genetic system to detect protein-protein interactions. Nature 340, 245–246. 10.1038/340245a0.

17. Koegl, M., and Uetz, P. (2007). Improving yeast two-hybrid screening systems. Brief Funct Genomic Proteomic 6, 302–312. 10.1093/bfgp/elm035.

18. Yu, H., Braun, P., Yildirim, M.A., Lemmens, I., Venkatesan, K., Sahalie, J., Hirozane-Kishikawa, T., Gebreab, F., Li, N., Simonis, N., et al. (2008). High-quality binary protein interaction map of the yeast interactome network. Science 322, 104–110. 10.1126/science.1158684.

19. Rolland, T., Taşan, M., Charloteaux, B., Pevzner, S.J., Zhong, Q., Sahni, N., Yi, S., Lemmens, I., Fontanillo, C., Mosca, R., et al. (2014). A proteome-scale map of the human interactome network. Cell 159, 1212–1226. 10.1016/j.cell.2014.10.050.

20. Luck, K., Kim, D.K., Lambourne, L., Spirohn, K., Begg, B.E., Bian, W., Brignall, R., Cafarelli, T., Campos-Laborie, F.J., Charloteaux, B., et al. (2020). A reference map of the human binary protein interactome. Nature 580, 402–408. 10.1038/s41586-020-2188-x.

21. Moscardó García, M., Aalto, A., Montanari, A.N., Skupin, A., and Gonçalves, J. (2025). Multi-omic network inference from time-series data. NPJ Syst Biol Appl 11, 114. 10.1038/s41540-025-00591-1.

22. Baião, A.R., Cai, Z., Poulos, R.C., Robinson, P.J., Reddel, R.R., Zhong, Q., Vinga, S., and Gonçalves, E. (2025). A technical review of multi-omics data integration methods: from classical statistical to deep generative approaches. Brief Bioinform 26. 10.1093/bib/bbaf355.

23. Warenda, A.J., Kauffman, S., Sherrill, T.P., Becker, J.M., and Konopka, J.B. (2003). Candida albicans septin mutants are defective for invasive growth and virulence. Infect Immun 71, 4045–4051. 10.1128/iai.71.7.4045-4051.2003.

24. Dagdas, Y.F., Yoshino, K., Dagdas, G., Ryder, L.S., Bielska, E., Steinberg, G., and Talbot, N.J. (2012). Septin-mediated plant cell invasion by the rice blast fungus, Magnaporthe oryzae. Science 336, 1590–1595. 10.1126/science.1222934.

25. Eisermann, I., Garduño-Rosales, M., and Talbot, N.J. (2023). The emerging role of septins in fungal pathogenesis. Cytoskeleton (Hoboken) 80, 242–253. 10.1002/cm.21765.

26. Li, L., Zhu, X.M., Su, Z.Z., Del Poeta, M., Liu, X.H., and Lin, F.C. (2021). Insights of roles played by septins in pathogenic fungi. Virulence 12, 1550–1562. 10.1080/21505594.2021.1933370.

27. Ryder, L.S., Cruz-Mireles, N., Molinari, C., Eisermann, I., Eseola, A.B., and Talbot, N.J. (2022). The appressorium at a glance. J Cell Sci 135. 10.1242/jcs.259857.

28. Shi, W., Cannon, K.S., Curtis, B.N., Edelmaier, C., Gladfelter, A.S., and Nazockdast, E. (2023). Curvature sensing as an emergent property of multiscale assembly of septins. Proc Natl Acad Sci U S A 120, e2208253120. 10.1073/pnas.2208253120.

29. Dulal, N., Rogers, A., Wang, Y., and Egan, M. (2020). Dynamic assembly of a higher-order septin structure during appressorium morphogenesis by the rice blast fungus. Fungal Genet Biol 140, 103385. 10.1016/j.fgb.2020.103385.

30. Howard, R.J., and Ferrari, M.A. (1989). Role of melanin in appressorium function. Experimental Mycology 13, 403–418. 10.1016/0147-5975(89)90036-4.

31. Sen, A., Mukherjee, D., and Aguilar, R.C. (2013). Bem3: Filling the GAP between cell polarity and secretion. Commun Integr Biol 6, e26702. 10.4161/cib.26702.

32. Costanzo, M., Baryshnikova, A., Bellay, J., Kim, Y., Spear, E.D., Sevier, C.S., Ding, H., Koh, J.L., Toufighi, K., Mostafavi, S., et al. (2010). The genetic landscape of a cell. Science 327, 425–431. 10.1126/science.1180823.

33. Fornerod, M., Ohno, M., Yoshida, M., and Mattaj, I.W. (1997). CRM1 is an export receptor for leucine-rich nuclear export signals. Cell 90, 1051–1060. 10.1016/s0092-8674(00)80371-2.

34. Costanzo, M., VanderSluis, B., Koch, E.N., Baryshnikova, A., Pons, C., Tan, G., Wang, W., Usaj, M., Hanchard, J., Lee, S.D., et al. (2016). A global genetic interaction network maps a wiring diagram of cellular function. Science 353. 10.1126/science.aaf1420.

35. Hecht, M., Rösler, R., Wiese, S., Johnsson, N., and Gronemeyer, T. (2019). An Interaction Network of the Human SEPT9 Established by Quantitative Mass Spectrometry. G3 (Bethesda) 9, 1869–1880. 10.1534/g3.119.400197.

36. Nakahira, M., Macedo, J.N., Seraphim, T.V., Cavalcante, N., Souza, T.A., Damalio, J.C., Reyes, L.F., Assmann, E.M., Alborghetti, M.R., Garratt, R.C., et al. (2010). A draft of the human septin interactome. PLoS One 5, e13799. 10.1371/journal.pone.0013799.

37. Foster, A., Were, V., Yan, X., Win, J., Harant, A., Langner, T., Kamoun, S., and Talbot, N. (2021). MagnaGenes (v. 1.0): an open science database of gene function studies in the blast fungus Magnaporthe oryzae. Zenodo. doi *10*.

38. Yan, X., Tang, B., Ryder, L.S., MacLean, D., Were, V.M., Eseola, A.B., Cruz-Mireles, N., Ma, W., Foster, A.J., Osés-Ruiz, M., and Talbot, N.J. (2023). The transcriptional landscape of plant infection by the rice blast fungus Magnaporthe oryzae reveals distinct families of temporally co-regulated and structurally conserved effectors. Plant Cell 35, 1360–1385. 10.1093/plcell/koad036.

39. Oliveira-Garcia, E., Yan, X., Oses-Ruiz, M., de Paula, S., and Talbot, N.J. (2024). Effector-triggered susceptibility by the rice blast fungus Magnaporthe oryzae. New Phytol 241, 1007–1020. 10.1111/nph.19446.

40. Querin, L., Sanvito, R., Magni, F., Busti, S., Van Dorsselaer, A., Alberghina, L., and Vanoni, M. (2008). Proteomic analysis of a nutritional shift-up in Saccharomyces cerevisiae identifies Gvp36 as a BAR-containing protein involved in vesicular traffic and nutritional adaptation. J Biol Chem 283, 4730–4743. 10.1074/jbc.M707787200.

41. Dulal, N., Rogers, A.M., Proko, R., Bieger, B.D., Liyanage, R., Krishnamurthi, V.R., Wang, Y., and Egan, M.J. (2021). Turgor-dependent and coronin-mediated F-actin dynamics drive septin disc-to-ring remodeling in the blast fungus Magnaporthe oryzae. J Cell Sci 134. 10.1242/jcs.251298.

42. Talbot, N.J. (2003). Functional genomics of plant-pathogen interactions. New Phytol 159, 1–4. 10.1046/j.1469-8137.2003.00809.x.

43. Gladfelter, A.S., Bose, I., Zyla, T.R., Bardes, E.S., and Lew, D.J. (2002). Septin ring assembly involves cycles of GTP loading and hydrolysis by Cdc42p. J Cell Biol 156, 315–326. 10.1083/jcb.200109062.

44. Sadian, Y., Gatsogiannis, C., Patasi, C., Hofnagel, O., Goody, R.S., Farkasovský, M., and Raunser, S. (2013). The role of Cdc42 and Gic1 in the regulation of septin filament formation and dissociation. Elife 2, e01085. 10.7554/eLife.01085.

45. Howard, R.J., Ferrari, M.A., Roach, D.H., and Money, N.P. (1991). Penetration of hard substrates by a fungus employing enormous turgor pressures. Proc Natl Acad Sci U S A 88, 11281–11284. 10.1073/pnas.88.24.11281.

46. de Jong, J.C., McCormack, B.J., Smirnoff, N., and Talbot, N.J. (1997). Glycerol generates turgor in rice blast. Nature 389, 244–244. 10.1038/38418.

47. Wilson, R.A., and Talbot, N.J. (2009). Under pressure: investigating the biology of plant infection by Magnaporthe oryzae. Nat Rev Microbiol 7, 185–195. 10.1038/nrmicro2032.

48. Leung, H., Borromeo, E.S., Bernardo, M.A., and Notteghem, J.L. (1988). Genetic analysis of virulence in the rice blast fungus Magnaporthe grisea. Phytopathology 78, 1227–1233.

49. Talbot, N.J., Ebbole, D.J., and Hamer, J.E. (1993). Identification and characterization of MPG1, a gene involved in pathogenicity from the rice blast fungus Magnaporthe grisea. Plant Cell 5, 1575–1590. 10.1105/tpc.5.11.1575.

50. Tingay, S., McElroy, D., Kalla, R., Fieg, S., Wang, M., Thornton, S., and Brettell, R. (1997). Agrobacterium tumefaciens-mediated barley transformation. The Plant Journal 11, 1369–1376.

51. Ryder, L.S., Dagdas, Y.F., Kershaw, M.J., Venkataraman, C., Madzvamuse, A., Yan, X., Cruz-Mireles, N., Soanes, D.M., Oses-Ruiz, M., and Styles, V. (2019). A sensor kinase controls turgor-driven plant infection by the rice blast fungus. Nature 574, 423–427.

52. Kershaw, M.J., and Talbot, N.J. (2009). Genome-wide functional analysis reveals that infection-associated fungal autophagy is necessary for rice blast disease. Proc Natl Acad Sci U S A 106, 15967–15972. 10.1073/pnas.0901477106.

53. Langmead, B., and Salzberg, S.L. (2012). Fast gapped-read alignment with Bowtie 2. Nat Methods 9, 357–359. 10.1038/nmeth.1923.

54. Li, H., Handsaker, B., Wysoker, A., Fennell, T., Ruan, J., Homer, N., Marth, G., Abecasis, G., and Durbin, R. (2009). The Sequence Alignment/Map format and SAMtools. Bioinformatics 25, 2078–2079. 10.1093/bioinformatics/btp352.

55. Danecek, P., Bonfield, J.K., Liddle, J., Marshall, J., Ohan, V., Pollard, M.O., Whitwham, A., Keane, T., McCarthy, S.A., Davies, R.M., and Li, H. (2021). Twelve years of SAMtools and BCFtools. Gigascience 10. 10.1093/gigascience/giab008.

56. Thorvaldsdóttir, H., Robinson, J.T., and Mesirov, J.P. (2013). Integrative Genomics Viewer (IGV): high-performance genomics data visualization and exploration. Brief Bioinform 14, 178–192. 10.1093/bib/bbs017.

57. Robinson, J.T., Thorvaldsdóttir, H., Winckler, W., Guttman, M., Lander, E.S., Getz, G., and Mesirov, J.P. (2011). Integrative genomics viewer. Nat Biotechnol 29, 24–26. 10.1038/nbt.1754.

58. Cox, J., and Mann, M. (2008). MaxQuant enables high peptide identification rates, individualized p.p.b.-range mass accuracies and proteome-wide protein quantification. Nat Biotechnol 26, 1367–1372. 10.1038/nbt.1511.

59. Yates, A.D., Allen, J., Amode, R.M., Azov, A.G., Barba, M., Becerra, A., Bhai, J., Campbell, L.I., Carbajo Martinez, M., Chakiachvili, M., et al. (2022). Ensembl Genomes 2022: an expanding genome resource for non-vertebrates. Nucleic Acids Res 50, D996–d1003. 10.1093/nar/gkab1007.

60. Cox, J., Neuhauser, N., Michalski, A., Scheltema, R.A., Olsen, J.V., and Mann, M. (2011). Andromeda: a peptide search engine integrated into the MaxQuant environment. J Proteome Res 10, 1794–1805. 10.1021/pr101065j.

61. Elias, J.E., and Gygi, S.P. (2010). Target-decoy search strategy for mass spectrometry-based proteomics. Methods Mol Biol 604, 55–71. 10.1007/978-1-60761-444-9_5.

62. Cox, J., Hein, M.Y., Luber, C.A., Paron, I., Nagaraj, N., and Mann, M. (2014). Accurate proteome-wide label-free quantification by delayed normalization and maximal peptide ratio extraction, termed MaxLFQ. Mol Cell Proteomics 13, 2513–2526. 10.1074/mcp.M113.031591.

63. Tyanova, S., Temu, T., and Cox, J. (2016). The MaxQuant computational platform for mass spectrometry-based shotgun proteomics. Nat Protoc 11, 2301–2319. 10.1038/nprot.2016.136.

64. Conway, J.R., Lex, A., and Gehlenborg, N. (2017). UpSetR: an R package for the visualization of intersecting sets and their properties. Bioinformatics 33, 2938–2940. 10.1093/bioinformatics/btx364.

65. Yon, G.G.V. (2021). Building, importing, and exporting gexf graph files with rgexf. Journal of Open Source Software 6, 3456.

66. Bastian, M., Heymann, S., and Jacomy, M. (2009). Gephi: An Open Source Software for Exploring and Manipulating Networks. Proceedings of the International AAAI Conference on Web and Social Media, 361-362. 10.1609/icwsm.v3i1.13937.

67. Jacomy, M., Venturini, T., Heymann, S., and Bastian, M. (2014). ForceAtlas2, a continuous graph layout algorithm for handy network visualization designed for the Gephi software. PLoS One 9, e98679. 10.1371/journal.pone.0098679.

68. Gu, Z., Eils, R., and Schlesner, M. (2016). Complex heatmaps reveal patterns and correlations in multidimensional genomic data. Bioinformatics 32, 2847–2849. 10.1093/bioinformatics/btw313.

69. Paysan-Lafosse, T., Blum, M., Chuguransky, S., Grego, T., Pinto, B.L., Salazar, G.A., Bileschi, M.L., Bork, P., Bridge, A., Colwell, L., et al. (2023). InterPro in 2022. Nucleic Acids Res 51, D418–d427. 10.1093/nar/gkac993.

70. Kinsella, R.J., Kähäri, A., Haider, S., Zamora, J., Proctor, G., Spudich, G., Almeida-King, J., Staines, D., Derwent, P., Kerhornou, A., et al. (2011). Ensembl BioMarts: a hub for data retrieval across taxonomic space. Database (Oxford) 2011, bar030. 10.1093/database/bar030.

71. Alexa, A., Rahnenführer, J., and Lengauer, T. (2006). Improved scoring of functional groups from gene expression data by decorrelating GO graph structure. Bioinformatics 22, 1600–1607. 10.1093/bioinformatics/btl140.

72. Benjamini, Y., and Hochberg, Y. (1995). Controlling the False Discovery Rate: A Practical and Powerful Approach to Multiple Testing. Journal of the Royal Statistical Society: Series B (Methodological) 57, 289–300. 10.1111/j.2517-6161.1995.tb02031.x.

73. Kolde, R. (2025). pheatmap Pretty Heatmaps. R package version 1.0. 12,(2018).

74. Cruz-Mireles, N., Osés-Ruiz, M., Derbyshire, P., Jégousse, C., Ryder, L.S., Bautista, M.J.A., Eseola, A., Sklenar, J., Tang, B., Yan, X., et al. (2024). The phosphorylation landscape of infection-related development by the rice blast fungus. Cell 187, 2557–2573.e2518. 10.1016/j.cell.2024.04.007.

75. Emms, D.M., and Kelly, S. (2019). OrthoFinder: phylogenetic orthology inference for comparative genomics. Genome Biol 20, 238. 10.1186/s13059-019-1832-y.

