## Supplementary figures and images for "A stage-resolved map of dynamic septin interactions required for infection by the rice blast fungus"

### Figure S1

#### 1179 Putative Septin Interactors Identified During Mycelium and Appressoirum Development

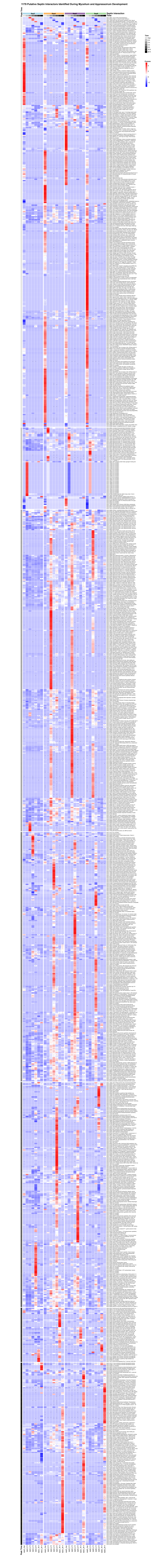

### Figure S2

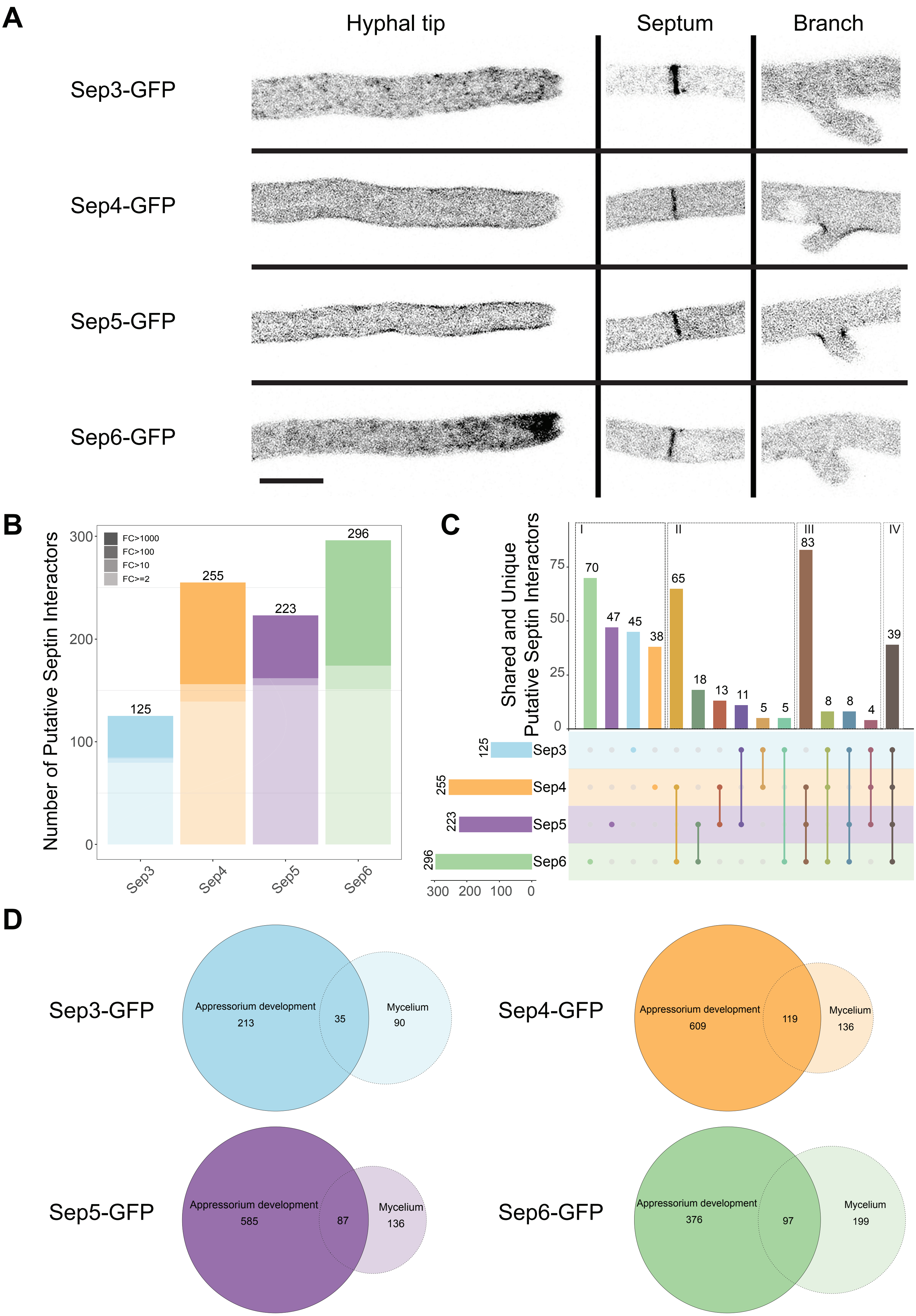

### Figure S3

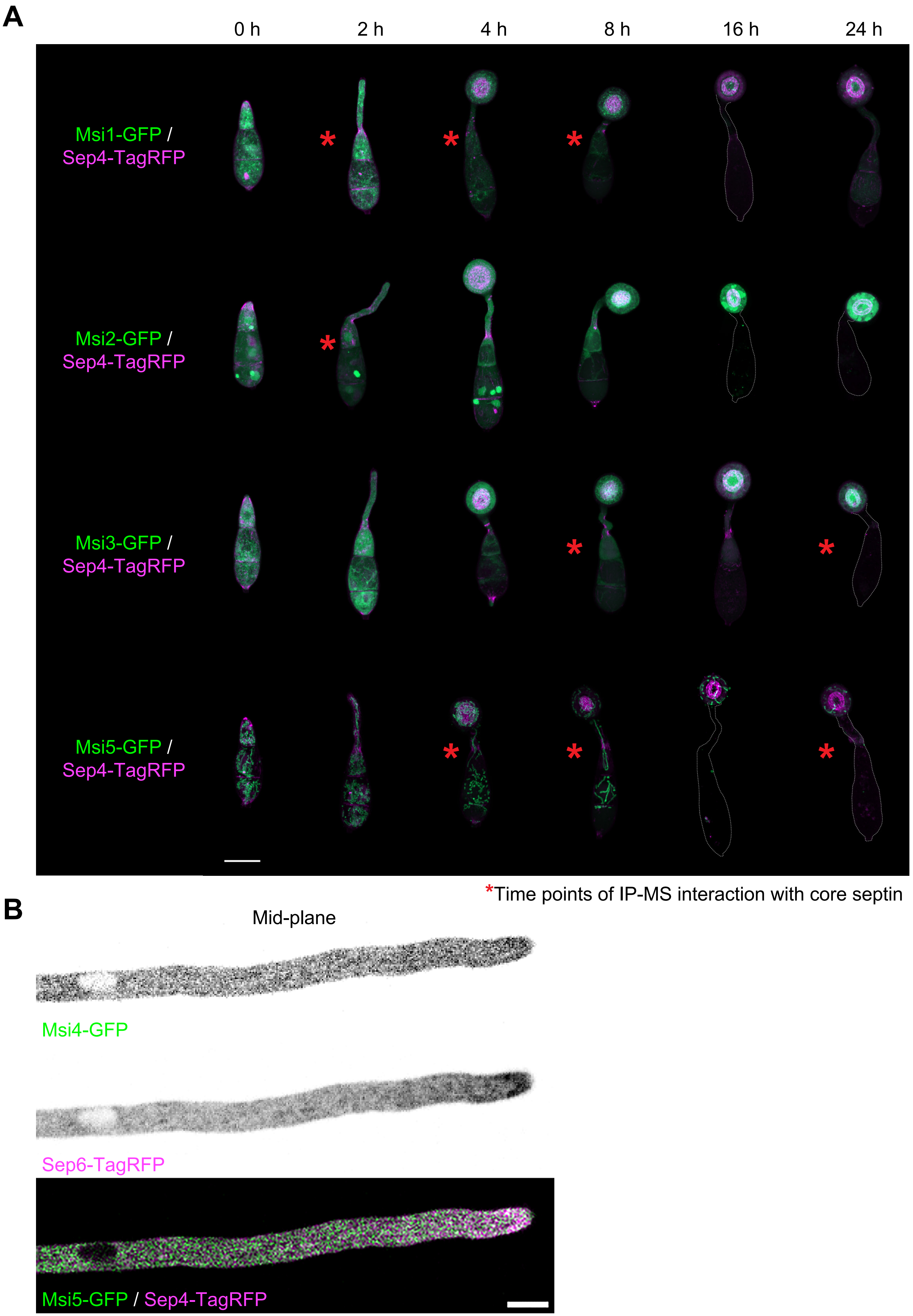

### Figure S4

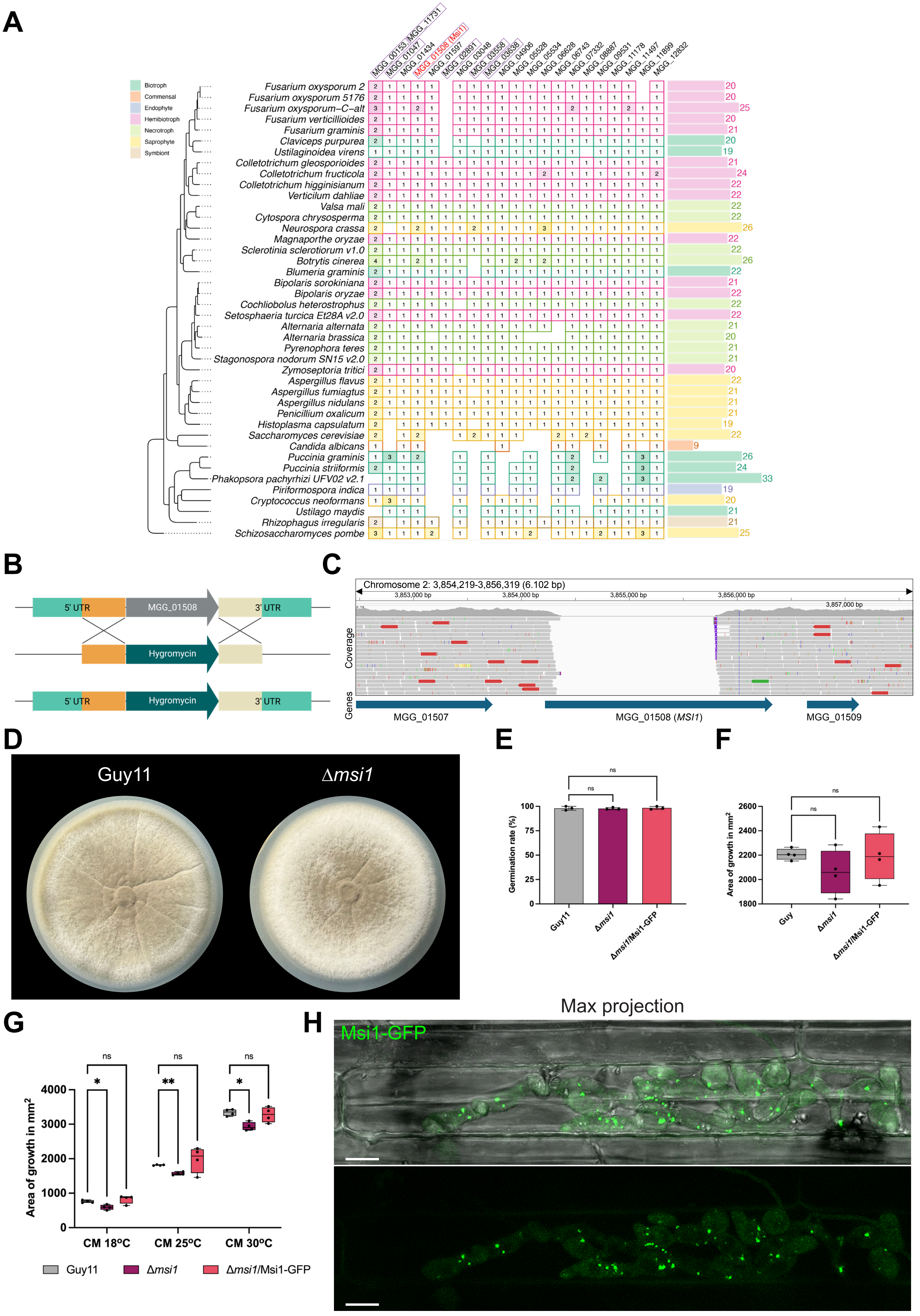
